# scDisInFact: disentangled learning for integration and prediction of multi-batch multi-condition single-cell RNA-sequencing data

**DOI:** 10.1101/2023.05.01.538975

**Authors:** Ziqi Zhang, Xinye Zhao, Peng Qiu, Xiuwei Zhang

## Abstract

Single-cell RNA-sequencing (scRNA-seq) has been widely used for disease studies, where sample batches are collected from donors under different conditions including demographical groups, disease stages, and drug treatments. It is worth noting that the differences among sample batches in such a study are a mixture of technical confounders caused by batch effect and the biological variations caused by condition effect. However, current batch effect removal methods often eliminate both technical batch effects and meaningful condition effects, while perturbation prediction methods solely focus on condition effects, resulting in inaccurate gene expression predictions due to unaccounted batch effects.

Here we introduce scDisInFact, a deep learning framework that models both batch effect and condition effect in scRNA-seq data. scDisInFact learns latent factors that disentangle condition effects from batch effects, enabling it to simultaneously perform three tasks: batch effect removal, condition-associated key gene detection, and perturbation prediction. We evaluated scDisInFact on both simulated and real datasets, and compared its performance to baseline methods for each task. Our results demonstrate that scDisInFact outperforms existing methods that focus on individual tasks, providing a more comprehensive and accurate approach for integrating and predicting multi-batch multi-condition single-cell RNA-sequencing data.

## 1 Introduction

Single-cell RNA-sequencing (scRNA-seq) is able to measure the expression levels of genes in each cell of an experimental batch. This technology has been widely used in disease studies, where samples are collected from donors at different stages of the disease or with different drug treatments^1–5^. As a result, each sample’s scRNA-seq count matrix is associated with one or more biological conditions of the donor, which can be age, gender, drug treatment, disease severity, etc. Meanwhile, datasets that study the same disease are often obtained in different batches, which introduce technical variations (also termed *batch effects*) across batches^6,7^. In practice, the available samples in the datasets of a disease study can originate from different conditions and batches, as arranged in Fig. 1a (right). We term such datasets as *multi-batch multi-condition datasets*. In such datasets, biological variations caused by *condition effects* exist between data matrices generated in the same batch but corresponding to different biological conditions, while technical variations caused by batch effects exist between data matrices from the same condition but different batches. Therefore, the differences among these data matrices are a mixture of batch effects (technical variation) and condition effects (biological variation), which complicates the process of fully utilizing the potential of these datasets.

**Figure 1.**
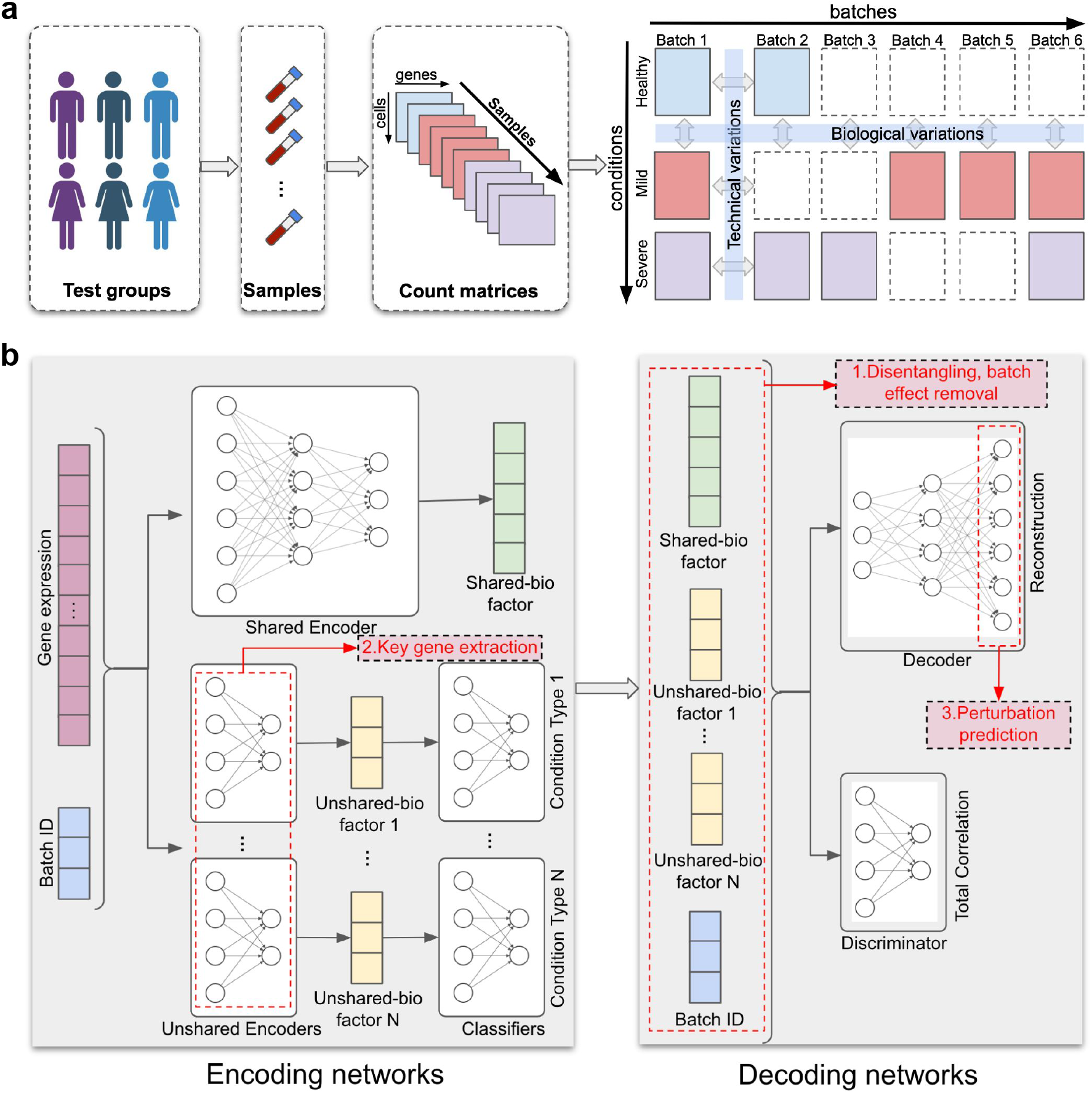
scDisInFact overview. **a**. Application scenarios of scDisInFact, where samples from disease study are organized into a multi-batch multi-condition dataset. **b**. The framework of scDisInFact, including an encoding neural network (left) and a decoding neural network (right).

In this paper, we consider a few computational challenges that need to be tackled in order to use multi-batch multi-condition datasets for disease study: (1) Removing batch effect while preserving biological condition effect; (2) Detecting key genes associated with biological conditions; (3) Predicting unseen data matrices corresponding to certain conditions (the matrices with dashed borders in Fig. 1a (right)), also known as the task of *perturbation prediction*. Methods have been designed for each problem separately, but no existing method can solve the three problems jointly. In the following, we discuss existing methods for each problem and their limitations.

Most existing batch effect removal methods treat the differences between data from different batches solely as batch effects and remove them by aligning different batches into a common distribution in either the original gene expression space or the latent embedding^8–12^. Applying these methods to data from multiple conditions can result in over-correction^13^, where biological differences among batches are also removed along with batch effects. Recently, methods have been proposed considering the biological differences among batches. scINSIGHT^14^ factorizes the scRNA-seq matrices into common and condition-specific modules using non-negative matrix factorization. However, the factorization framework is limited to one type of condition, while multiple types of conditions, such as age, gender, and drug treatment, can co-exist in the dataset^15,16^. In addition, scINSIGHT corrects the batch effect on the latent space and cannot predict gene expression matrices that are removed of the batch effect under various conditions. scMC^13^ integrates data batches without removing the biological variations among batches, but it does not disentangle biological variations caused by condition effect from those that are shared among batches, nor does it output key genes associated with the biological conditions.

Another problem in the field is predicting the scRNA-seq data under one condition using data from another condition, also known as the problem of *perturbation prediction*^17^. This is particularly useful when predicting disease progression or drug effects, under conditions where data are not collected. Existing methods for this task, such as scGen^17^ and scPreGAN^18^, do not account for batch effects between data matrices and assume differences in cell distribution between data matrices solely result from biological conditions, an assumption that does not hold for most real datasets. Furthermore, in practice, there is often more than one type of condition in the data, but existing methods are designed for only one type of condition. For example, conditions such as disease severity and treatment can exist at the same time in a disease study, but existing methods can not deal with both types of conditions at the same time.

Here, we propose scDisInFact (**s**ingle **c**ell **dis**entangled **In**tegration preserving condition-specific **Fact**ors), which is the first method that can perform all three tasks: batch effect removal, condition-associated key genes (CKGs) detection, and perturbation prediction on multi-batch multi-condition scRNA-seq dataset (Fig. 1a). scDisInFact is designed based on a disentangle variational autoencoder framework. It disentangles the variation within the multi-batch multi-condition dataset into latent factors encoding the biological information shared across all data matrices, condition-specific biological information, and technical batch effect. The disentangled latent space allows scDisInFact to perform two other tasks, the CKG detection and perturbation prediction, and to overcome the limitation of existing methods for each task. In particular, the disentangled factors allow scDisInFact to remove batch effect while keeping the condition effect in gene expression data. In addition, scDisInFact expands the versatility of existing perturbation prediction methods in that (1) it models the effect of multiple condition types and (2) it enables the prediction of data across any combination of conditions and batches within the dataset.

We compared scDisInFact with scINSIGHT in terms of batch effect removal and CKG detection. As scINSIGHT does not perform perturbation prediction, we compared scDisInFact with scGen and scPreGAN in terms of perturbation prediction. We tested scDisInFact on simulated and real datasets^1–4^, and found that it outperforms baseline methods across various tasks. Owing to its superior performance and multi-task capabilities, scDisInFact can be employed to comprehensively analyze multi-batch multi-condition scRNA-seq datasets, facilitating a deeper understanding of disease progression and patient responses to drug treatments.

## 2 Results

### 2.1 Overview of scDisInFact

scDisInFact is designed for a general multi-batch multi-condition scRNA-seq dataset scenario where samples are obtained from donors of different conditions and profiled in multiple experimental batches (Fig. 1a). We termed the category of conditions as the *condition type*, and the condition label of each condition type as a *condition*. For example, given a dataset that is obtained from donors with varying disease severity and genders, disease severity and gender are considered two separate *condition types*, while specific severity levels such as healthy control, mild symptoms, and severe symptoms are *conditions*. When there are more than one condition types, a *condition* can be the combination of labels, each label from a condition type. For each cell, scDisInFact takes as input not only its gene expression data, but also its batch ID and identified condition of each condition type.

scDisInFact is designed based on a variational autoencoder (VAE) framework^19,20^ (Fig. 1b). In the model, the encoder networks encode the high dimensional gene expression data of each cell into a disentangled set of latent factors, and the decoder network reconstructs gene expression data from the latent factors (Fig. 1b). scDisInFact has multiple encoder networks, where each encoder learns independent latent factors from the data. Using the information of biological condition and batch ID of each cell, scDisInFact effectively disentangles the gene expression data into the shared biological factors (*shared-bio factors*), unshared biological factors (*unshared-bio factors*), and technical batch effect.

The shared encoder learns shared-bio factors (green vector in Fig. 1b), which represent the biological variations within a data matrix that are irrelevant to condition effect or batch effects. Such variations usually are reflected as heterogeneous cell types in individual data matrices. An unshared encoder learns unshared-bio factors, which represent biological variations that are related to the condition effect. The number of unshared encoders matches the number of condition types, with each unshared encoder learning unshared-bio factors exclusively for its corresponding condition type (yellow vectors in Fig. 1b). For example, given a dataset of donors with different disease severities and genders, two unshared encoders are used, where one learns the disease severity condition and the other learns the gender condition. The technical batch effect is encoded as pre-defined one-hot batch factors (blue vector in Fig. 1b) which are transformed from the batch IDs of each cell. The batch factors are appended to the gene expression data before being fed into each encoder (Fig. 1b (left)) in order to differentiate the gene expression data of cells from different batches. The decoder takes as input the shared-bio factors, unshared-bio factors, and batch factors, and reconstructs the input gene expression data (Fig. 1b (right)). In order to guide the unshared encoders to extract biological variations that are only relevant to condition effects, a linear classifier network is attached to the output of each unshared encoder, and is trained together with the unshared encoder to predict the conditions of the cells (Fig. 1b (left)). A discriminator network is used for disentangling the shared-bio and unshared-bio factors, following the approach of Factor-VAE^21^ (Fig. 1b (right)).

With the disentangled latent factors, scDisInFact can perform the three tasks it promises (red text in pink boxes in Fig. 1b): (1) Batch effect removal. The learned shared-bio factors are removed of the batch effect and condition effect and can be used as cell embedding for clustering and cell type identification. Combining the shared-bio factors and any unshared-bio factors gives cell embeddings including the corresponding condition effects. (2) Detection of condition-associated key genes (CKGs) for each condition type. The first layer of each unshared encoder is designed to be a key feature (gene) selection layer of the corresponding condition type. The final weights of that layer can be transformed into the gene scores associated with the condition type (Methods, Fig. S1a). (3) Perturbation prediction. After scDisInFact is trained and the shared-bio factors, unshared-bio factors, and batch factors are learned, the perturbation prediction task takes the gene expression data of a set of cells under a certain condition and batch as input, and predicts the gene expression data of the same cells under a different condition in the same or different batch (Fig. S1b). Fig. S1b shows how to predict data in condition 2 and batch 1 given data in condition 1 and batch 6. We first calculate new unshared-bio factors and batch factors according to the predicted condition. The decoder then generate the predicted data matrix using the new unshared bio-factors and batch factors (Methods). It is worth noting that scDisInFact can perform perturbation prediction from any condition and batch to any other condition and batch in the dataset (Methods).

The loss formulation, training procedure, and procedures to obtain the results for each task from a trained scDisInFact model are included in Methods.

### 2.2 Tests on simulated dataset

We first validate scDisInFact on simulated datasets where we have ground truth information. We generated simulated datasets with 2 batches and 2 condition types including treatment and disease severity. The first condition type (treatment) includes cells under control (*ctrl*) and stimulation (*stim*) conditions, and the second condition type (severity) includes *healthy* and *severe* conditions. The condition and batch arrangement of count matrices in the dataset follows Fig. 2a. Totally 9 datasets were generated with different numbers of CKGs and perturbation parameters, which models different strengths of condition effect (Methods, parameter settings of each dataset follow Table S1). In the following, we first show the generalization ability of scDisInFact, then show the performance of scDisInFact in terms of the three tasks, batch effect removal (which is also latent space disentanglement), key gene detection, and perturbation prediction.

**Figure 2.**
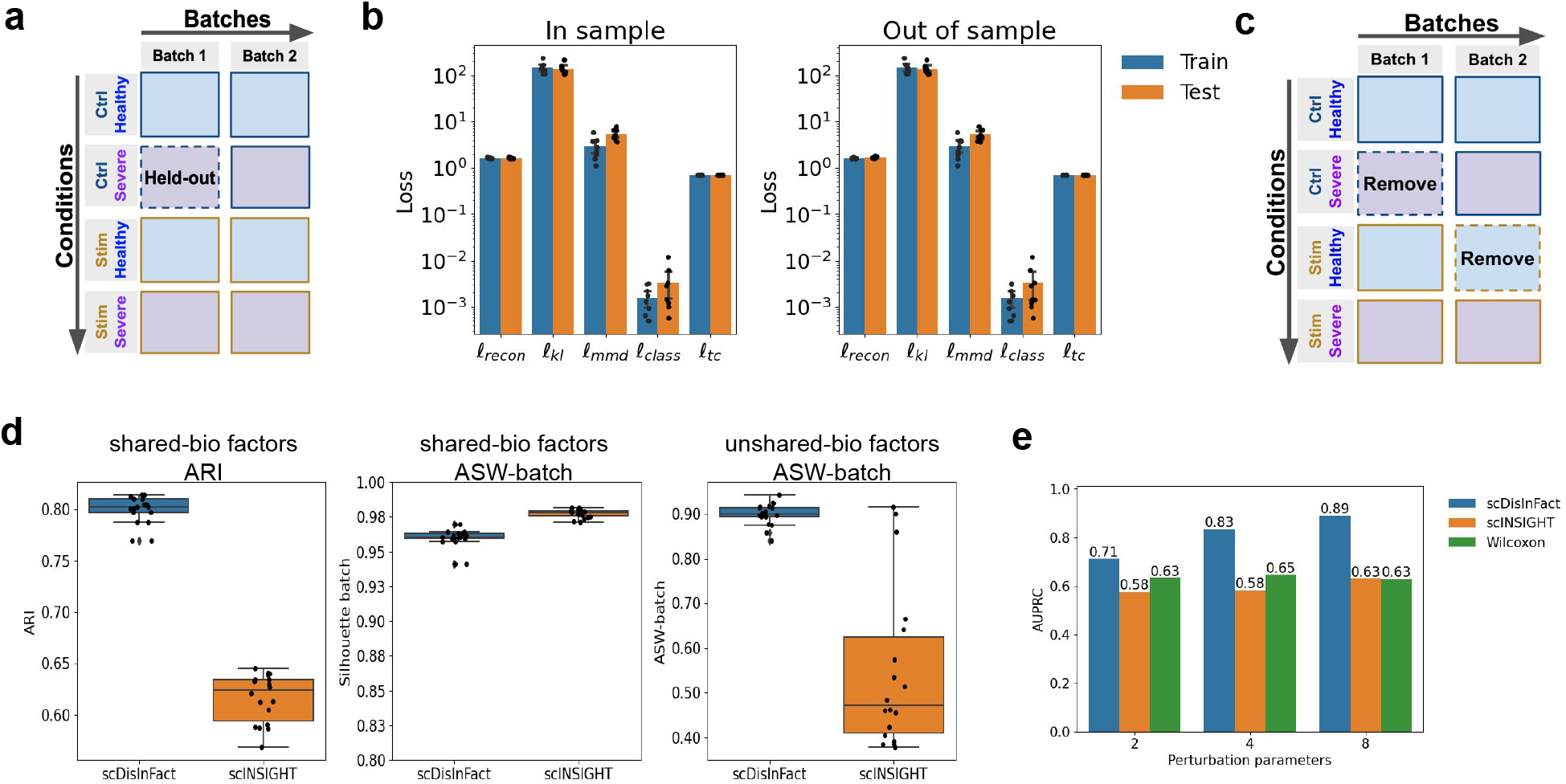
Results on simulated datasets. **a**. The arrangement of count matrices in the simulated datasets. The matrix with a dashed border is held out in the out-of-sample generalization test. **b**. The loss values of the model in training and testing datasets, including both in-sample test (left) and out-of-sample test (right). **c**. The arrangement of count matrix in disentanglement test. The matrices with dashed borders are removed in the test. **d**. The disentanglement scores of scDisInFact and scINSIGHT. **e**. AUPRC of scDisInFact, scINSIGHT, and Wilcoxon rank sum test on CKGs detection.

#### 2.2.1 Test generalization ability of scDisInFact

One potential problem with neural network models is overfitting. If the model overfits, the model cannot be generalized and cannot be used for prediction tasks. Therefore, generalization ability is the basic requirement of scDisInFact in order to achieve successful disentanglement and make accurate perturbation predictions. We test the generalization ability of scDisInFact using the simulated dataset with 2 condition types (Fig. 2a).

We test the generalization ability of scDisInFact under two different scenarios, according to the relationship between training and testing datasets, referred to as *in-sample* test and *out-of-sample* test. In the *in-sample* test, the training and testing data are evenly sampled from the same original dataset and they follow the same data distribution. We randomly held out 10% of cells in each count matrix as test data, and trained the model on the remaining 90% of cells. In the *out-of-sample* test, the training and testing data are not evenly sampled from the original dataset and the test distribution no longer match the training distribution. We held out the count matrix corresponding to the condition <*ctrl*, *severe*> in batch 1, and trained the model using the remaining count matrices (Fig. 2a, held-out matrix shown with a dashed border). We expect that it is harder for the model to generalize for the “out-of-sample” test due to the distribution difference.

After the model is trained on the training data, we tested it on the held-out data and compared the training and testing losses. The loss values on the 9 simulated datasets (Fig. 2b) show that in both scenarios (in-sample test and out-of-sample test), the testing losses are close to the training losses for various loss terms, confirming the generalization ability of scDisInFact.

#### 2.2.2 scDisInFact performs latent space disentanglement and batch effect removal on simulated datasets

In this section, we tested the latent space disentanglement of scDisInFact on the simulated datasets, and compared its performance with scINSIGHT^14^. In practice, each batch often includes only a subset of the condition combinations. Therefore, to make the test case more realistic, we remove the count matrices corresponding to the <*ctrl*, *severe*> condition in batch 1 and the <*stim*, *healthy*> condition in batch 2 (Fig. 2c, removed matrices shown with dash borders). Since scINSIGHT can only work with one condition type, we ran scINSIGHT separately for each condition type (while fixing the other condition type): to learn the unshared bio-factor of *ctrl* and *stim* conditions, we fix the severity condition to *healthy* and train scINSIGHT using the 3 count matrices corresponding to <*ctrl*, *healthy*> and <*stim*, *healthy*> conditions. To learn the unshared bio-factor of *healthy* and *severe* conditions, we fix treatment to *ctrl* and train scINSIGHT on the 3 count matrices corresponding to <*ctrl*, *healthy*> and <*ctrl*, *severe*> conditions. The calculation of shared-bio and unshared-bio factors for scINSIGHT is described in Methods.

A successful disentanglement requires that: (1) The shared-bio factors encode the cell-to-cell biological variations irrelevant to condition and batch effect. With these factors, we expect cells should be grouped according to cell types and aligned across batches and conditions. (2) The unshared-bio factors encode condition-specific biological variation across data matrices irrelevant to batch effect. With these factors, cells should be grouped according to conditions, and aligned across batches under each condition.

We evaluate the learned shared-bio factor in terms of batch effect removal (using ASW-batch score, Methods) and separation of cell types (using ARI score, Methods). The boxplots of these scores (Fig 2d, left and middle) show that scDisInFact has a significantly better performance in separating cell types while removing the batch effect compared to scINSIGHT. When evaluating the unshared-bio factors, we measure the grouping of conditions using ARI score, and the removal of batch effect using ASW-batch score (Methods). Both methods have perfect ARI scores (equal to 1, plots not shown), which is expected as both methods enforce the grouping in their objective function. In the meantime, scDisInFact achieves a higher ASW-batch score, and shows a better performance in removing the batch effect in the unshared-bio factors (Fig. 2d, right).

To visually assess the disentanglement results, we visualized the learned shared-bio factors and unshared-bio factors from one simulated dataset (Simulation 1 in Table S1). We visualized the shared-bio factors using UMAP and observe that cell types are well-separated and batches are well-aligned (Fig. S2a). We visualized the unshared-bio factors corresponding to each condition type using PCA, and notice that the unshared-bio factors for each condition type effectively separate the conditions within that type, while the batches are still aligned (Fig. S2b,c).

#### 2.2.3 scDisInFact detects condition-associated key genes

Using the training result from the previous section, we further compared the CKG detection accuracy of scDisInFact and scINSIGHT. We also included Wilcoxon rank sum test as an additional baseline method. For each method, we obtain a CKG score for each gene to indicate how likely this gene is a CKG. For scINSIGHT, we used the variance of gene membership matrices across conditions as CKG scores (Methods). In Wilcoxon rank sum test, for each gene, we ran the test between the gene’s expression levels under different conditions, and obtained the corresponding *p*-values (with false discovery rate multi-tests correction). We transformed the *p*-values into CKG scores following 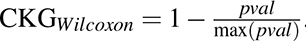.

AUPRC scores were calculated for each method between the inferred CKG scores and ground truth CKGs (Methods). We aggregate the mean AUPRC score of each method on the simulated datasets with different perturbation parameters *ε* in a barplot (Fig. 2e). scDisInFact consistently performs better than the two baseline methods under all values of the perturbation parameter *ε*, which shows that the successful disentanglement of biological variations and technical batch effect could help to better uncover the CKGs. The AUPRC score increases with the increase of the perturbation parameter. This is because a higher perturbation parameter corresponds to a larger difference in the expression of the CKGs across conditions, which makes it easier for the CKGs to be detected.

#### 2.2.4 scDisInFact performs accurate perturbation prediction under various scenarios

We test the perturbation prediction accuracy of scDisInFact on the same set of simulated datasets (Fig. 2a). Similar to the generalization test, we conducted perturbation prediction under two different scenarios: (1) In-sample prediction, where the condition to predict is seen in the training dataset, and (2) Out-of-sample prediction, where the condition to predict is not seen in the training dataset.

In the in-sample test, we train scDisInFact using all count matrices in Fig. 2a, and take one count matrix as input to predict the mRNA counts of the same cells under a different conditions. For example, in Fig. S3a, arrow #2 means that from the data matrix *X*_11_, predict the cells’ expression levels under batch 1, condition <*control*, *severe*>, denoted by 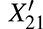. We use the notation 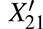 to distinguish the predicted matrix from *X*_21_ which is part of the training data. 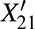 and *X*_21_ are matrices under the same batch and conditions but on different cells. Therefore, when evaluating the accuracy of 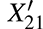, it can not be compared with *X*_21_ at the single cell level, but at the cell type level instead (Methods).

In the out-of-sample test, we held out all count matrices under condition <*ctrl*, *severe*> (*X*_21_*, X*_22_ in Fig. S3b), and trained the model on the remaining count matrices. Then we took one count matrix as input and predicted the corresponding counts under the held-out condition. In the out-of-sample test, <*ctrl*, *severe*> is the unseen condition because the combination of *ctrl* and *severe* is not seen in the training dataset, although the *ctrl* or *severe* condition can appear in the training data in other condition combinations.

For both the in-sample and out-of-sample predictions, we predict data under condition <*ctrl*, *severe*> using different matrices as input (Fig. S3a-b). Depending on which effects exist between the input and predicted matrices, we categorized the prediction test into 6 scenarios. When the input and predicted matrices are from the same batch, we test the prediction of condition effect of (1) condition Type 1 (treatment, arrow #1 in Fig. S3a,b), (2) condition Type 2 (severity, arrow #2 in Fig. S3a,b) and (3) condition Types 1 & 2 (arrow #3 in Fig. S3a,b). Similarly, when the input and predicted matrices are from different batches, we also test the prediction of condition effect of (4) condition type 1 (arrow #4 in Fig. S3a,b), (5) condition type 2 (arrow #5 in Fig. S3a,b) and (6) condition type 1 & 2 (arrow #6 in Fig. S3a,b). scDisInFact can predict data across any condition and batch combinations, allowing for the prediction of gene expression data for all the cells of all given data matrices under any condition and batch combination.

We compare the performance of scDisInFact with scGen and scPreGAN. As scGen and scPreGAN are only designed for one condition type, we train the methods using the count matrices with fixed condition values for the condition types that we are not predicting (Methods). The detailed settings of the training, input, and predicted data matrices for all methods are in Tables S2-3. We take the known count matrix under the predicted condition and batch, and use this matrix or its denoised matrix as the gold-standard counts (Methods). We compare the predicted counts with the gold-standard counts using cell-type-specific MSE, *R*^2^, and Pearson correlation scores (Methods). The results are summarized in Fig. 3a (in-sample tests) and Fig. 3b (out-of-sample tests). scDisInFact has the smallest MSE and highest Pearson correlation and *R*^2^ out of all three methods in all test scenarios. The baseline methods show much higher MSE (note that the y-axis uses log scale in the MSE plots) than scDisInFact while their Pearson correlation can be close to that of scDisInFact. The high MSE of baseline methods is caused by the difference in the distribution between predicted values and ground truth values, although normalization has been performed (Methods).

**Figure 3.**
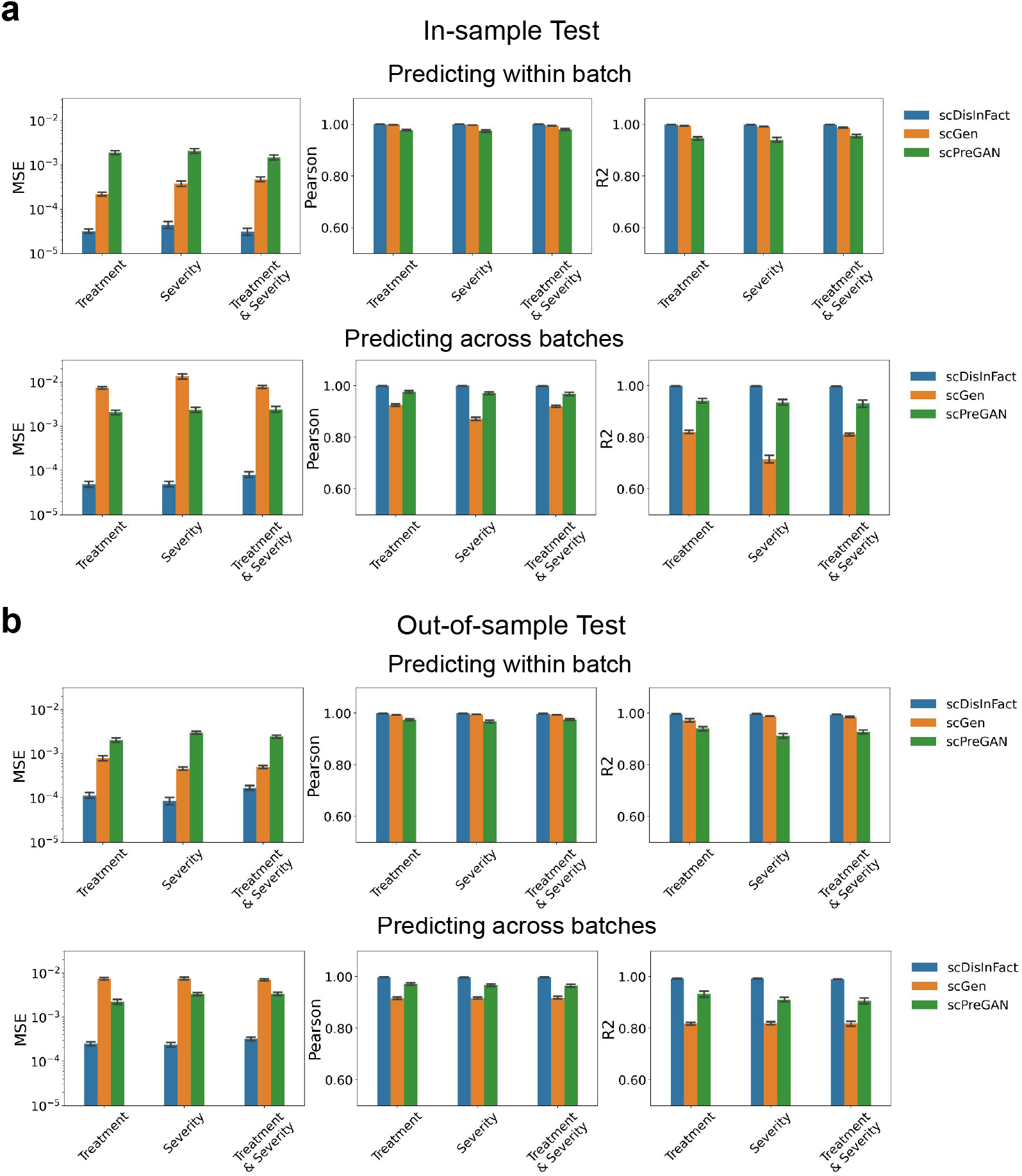
Perturbation prediction result on simulated datasets. **a**. Barplots showing the perturbation prediction accuracy (measured in cluster-specific MSE, Pearson correlation, and *R*^2^ score) of scDisInFact and baseline methods in in-sample test. The upper row shows the result where the input and predict count matrices are from the same batch, and the lower row shows the result where the input and predict count matrices are from different batches. (b) Barplot showing the perturbation prediction accuracy of scDisInFact and baseline methods in out-of-sample test. Plots are organized following **a**.

Since scDisInFact learns the condition and batch effects from the training dataset, the training dataset with smaller coverage of possible conditions and batches should affect the performance of scDisInFact. We further conducted experiments to analyze how the number of held-out matrices affects the prediction power of scDisInFact. We created 4 scenarios by holding out 1 to 4 count matrices from the training set. The detailed settings of the test scenarios are summarized in Table S4. After training the model, we take as input the count matrix corresponding to condition <*stim*, *severe*> in batch 2 (matrix *X*_42_ in Fig. S3c), and predict the counts under condition <*ctrl*, *severe*> of batch 1 (matrix 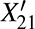 in Fig. S3c). We measure the cell-type-specific MSE, *R*^2^, and Pearson correlation scores (Methods) between the predicted and gold-standard counts (matrices *X*_21_ and 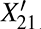) and aggregate the scores into the boxplot in Fig. S3d. From Fig. S3d, we observe the overall performance of scDisInFact drops when fewer matrices are included in the training data. However, even when holding out 3 matrices, the performance of scDisInFact is comparable or better than the better method out of scGen and scPreGAN for each metric in Fig. 3. With 4 matrices held out (Fig. S3c), the performance of scDisInFact deteriorates, as in this case, the difference between any two matrices is a mixture of condition effect and batch effect, which poses a challenging case for disentanglement.

### 2.3 Testing scDisInFact on glioblastoma dataset

We then applied scDisInFact to real datasets. We first applied scDisInFact on a glioblastoma (GBM) dataset^1^. The dataset has 21 count matrices from 6 patient batches with 1 condition type (drug treatment) that includes conditions: no-drug control (*vehicle (DMSO)*), and panobinostat drug treatment (*0.2 uM panobinostat*). The metadata of the count matrices follows Table S5 and the data matrices can be arranged in a “condition by batch” grid as shown in Fig. S4a. Before applying scDisInFact, we filtered the lowly-expressed genes (Methods) and visualized the dataset using UMAP. Strong technical batch effect and condition effect can be observed among batches and conditions (Fig. S4b).

After applying scDisInFact to pre-processed data, we obtained the shared-bio factors and unshared-bio factors that are specific to the two conditions (Figs. 4a, S4c). The shared-bio factors separate cells of major cell types (Fig. 4a (left)) and are removed of the batch and condition effect (Figs. 4a (middle, right)). The unshared-bio factors separate cells into different conditions (Fig. S4c (left)), and are removed of the batch effect (Fig. S4c (right)). The latent space visualization shows that scDisInFact is able to disentangle latent factors and remove the strong batch effects in the dataset. We compared the disentanglement results of scDisInFact and scINSIGHT both visually and quantitatively. The calculation of shared-bio and unshared-bio factors of scINSIGHT is described in Methods. We visualized the shared-bio and unshared-bio factors of scINSIGHT using UMAP (Fig. S5a). From Fig. S5a, the cell types are not well separated in the shared-bio factors (top-left plot) and the batches are not well aligned in unshared-bio factors (bottom-right plot). To quantitatively verify this, we evaluate the shared-bio factor using ASW-batch score and ARI score, where ASW-batch measures the removal of batch effect and ARI measures the separation of cell types (Methods, Fig. S5b). We evaluate the removal of batch effect in unshared-bio factor using ASW-batch scores (Fig. S5b). scDisInFact largely outperforms scINSIGHT in terms of both ARI score for shared-bio factors and ASW-batch score for unshared-bio factors, which is consistent with the visualizations.

**Figure 4.**
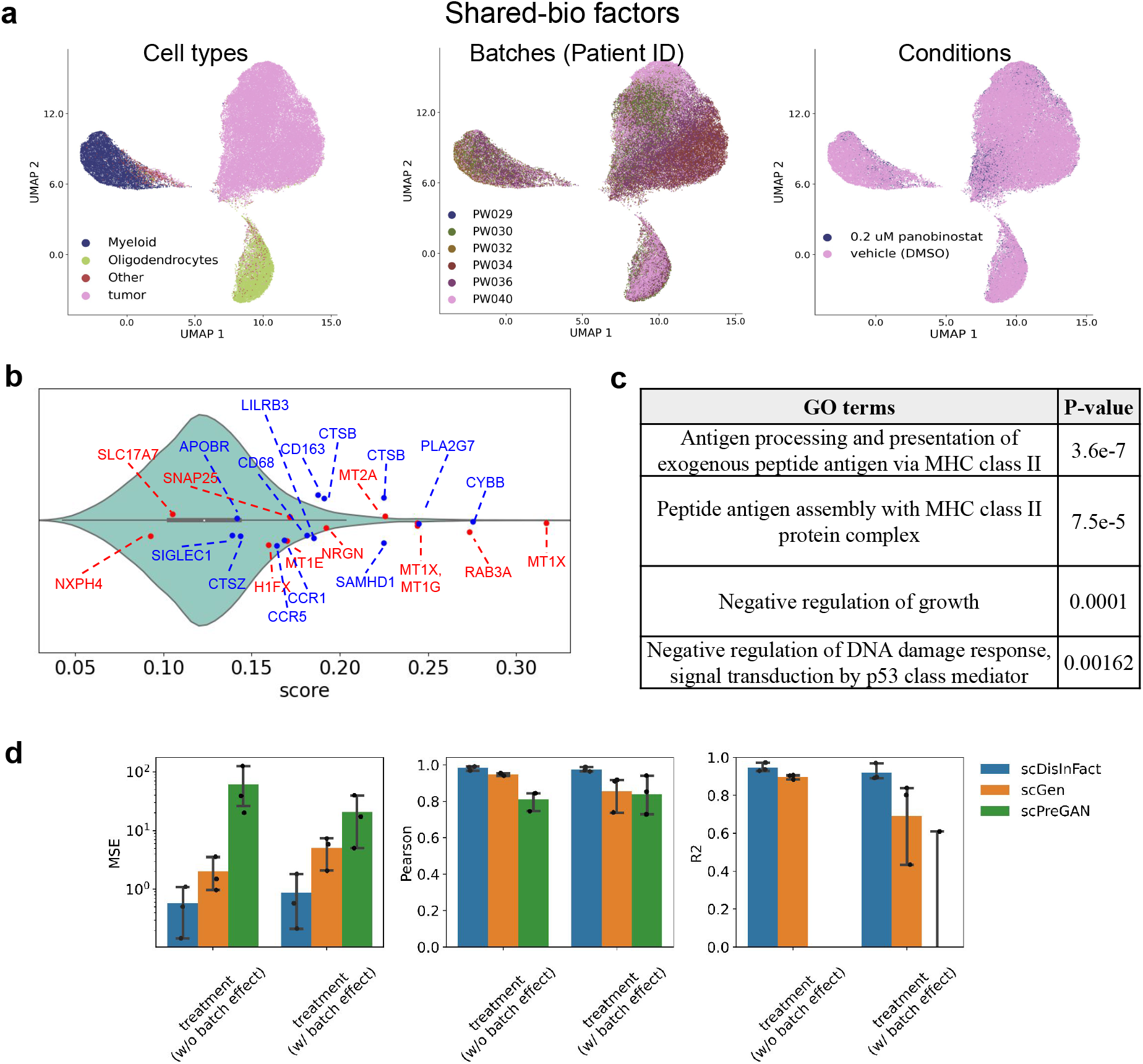
Test results on GBM dataset. **a**. UMAP visualization of shared-bio factors, where the cells are colored by (left) original cell type, (middle) batches, and (right) conditions. **b**. Violin plot of the CKG scores. Red dots correspond to metallothioneins and neuronal marker genes, and blue dots correspond to macrophage marker genes. **c**. Top GO terms inferred from top-scoring genes. **d**. Barplots showing the perturbation prediction accuracy of scDisInFact and baseline methods, where the cell-type-specific MSE, Pearson, and *R*^2^ scores are included.

We then analyzed the CKGs detected by scDisInFact. After training the model, we obtained the CKG score of each gene from the unshared encoder, and sorted the genes by their scores (Methods, Fig. 4b). We expect that the top-scoring genes are related to the biological processes associated with panobinostat drug treatment. We obtained high CKG scores for metallothioneins and neuronal marker genes (red dots in Fig. 4b), and macrophage marker genes (blue dots in Fig. 4b)^1^. These top-scoring marker genes are closely related to panobinostat treatment because panobinostat was reported to up-regulate the metallothionein family genes and mature neuronal genes, and down-regulate macrophage marker genes which were immunosuppressive in GBM^1^. We further conducted gene ontology (GO) analysis on the top-300 scoring genes using TopGO^22^, and the result shows multiple biological processes relevant to the panobinostat treatment in cancer (Fig. 4c). Panobinostat was shown to activate MHC II pathways^23^ and we do observe two terms that are related to MHC class II pathway proteins. There are also terms related to p53 transcription factor and regulation of growth. P53 is a tumor suppressor involved in the regulation of DNA damage response, which is affected by the use of panobinostat^24^. Panobinostat also was shown to have a growth suppressive effect on cancer^25^ (Fig. 4c).

We also tested the perturbation prediction accuracy of scDisInFact on the GBM dataset. We held out the count matrix corresponding to sample “PW034-705” (Table S5, Fig. S4a), and train scDisInFact on the remaining count matrices in the dataset. After training the model, we take one count matrix in the training dataset as input, and predict the counts for the same cells under the condition and batch of the held-out matrix. We choose different count matrices as input to create different scenarios for the prediction task: (1) the input and held-out matrices are from the same batch but under different conditions, where scDisInFact predicts only the condition effect; (2) the input and held-out matrices are from different batches and different conditions, where scDisInFact predicts both the condition effect and batch effect. The configurations of input and output data matrices are in Table S6.

We compared the performance of scDisInFact with scGen and scPreGAN for these prediction tasks using cell-type-specific MSE, Pearson correlation, and *R*^2^ scores (Methods) between the predicted and held-out (gold-standard) counts (Methods). We calculate the scores for cell types including “Myeloid”, “Oligodendrocytes”, and “tumor”, and aggregate the scores in Fig. 4d. From the comparison result, we observe that scDisInFact has a better prediction accuracy, and the prediction improvement is more pronounced when the batch effect exists between the input and held-out count. This is because scDisInFact specifically models batch effect in the perturbation prediction while the other two methods do not. In both prediction tasks, we further jointly visualize the predicted and the held-out counts using UMAP (Fig. S6a for within-batch prediction, Fig. S6b for cross-batch prediction). The visualization (Figs. S6a,b) shows that the predicted counts of scDisInFact match the held-out counts in both tasks while both baseline methods fail especially in the second task. The visualization result matches the quantitative measurement in Fig. 4d.

### 2.4 scDisInFact performs disentanglement and predicts data under unseen conditions on COVID-19 dataset

We then tested scDisInFact on a COVID-19 dataset with multiple condition types. We built a COVID-19 dataset by collecting data from three different studies^2–4^. Since no significant batch effect is reported within each study^2–4^, we treated the dataset obtained from each study as one batch. The resulting dataset has 3 batches of cells with 2 condition types: age and disease severity. The three batches are termed “Arunachalam_2020”, “Lee_2020”, and “Wilk_2020” according to the source of the study. The age condition includes young (*40-*), middle-aged (*40-65*), and senior (*65+*) groups, and the disease severity condition includes *healthy control*, *moderate symptom*, and *severe symptom* (Methods, the arrangement of count matrices follows Fig. 5a).

**Figure 5.**
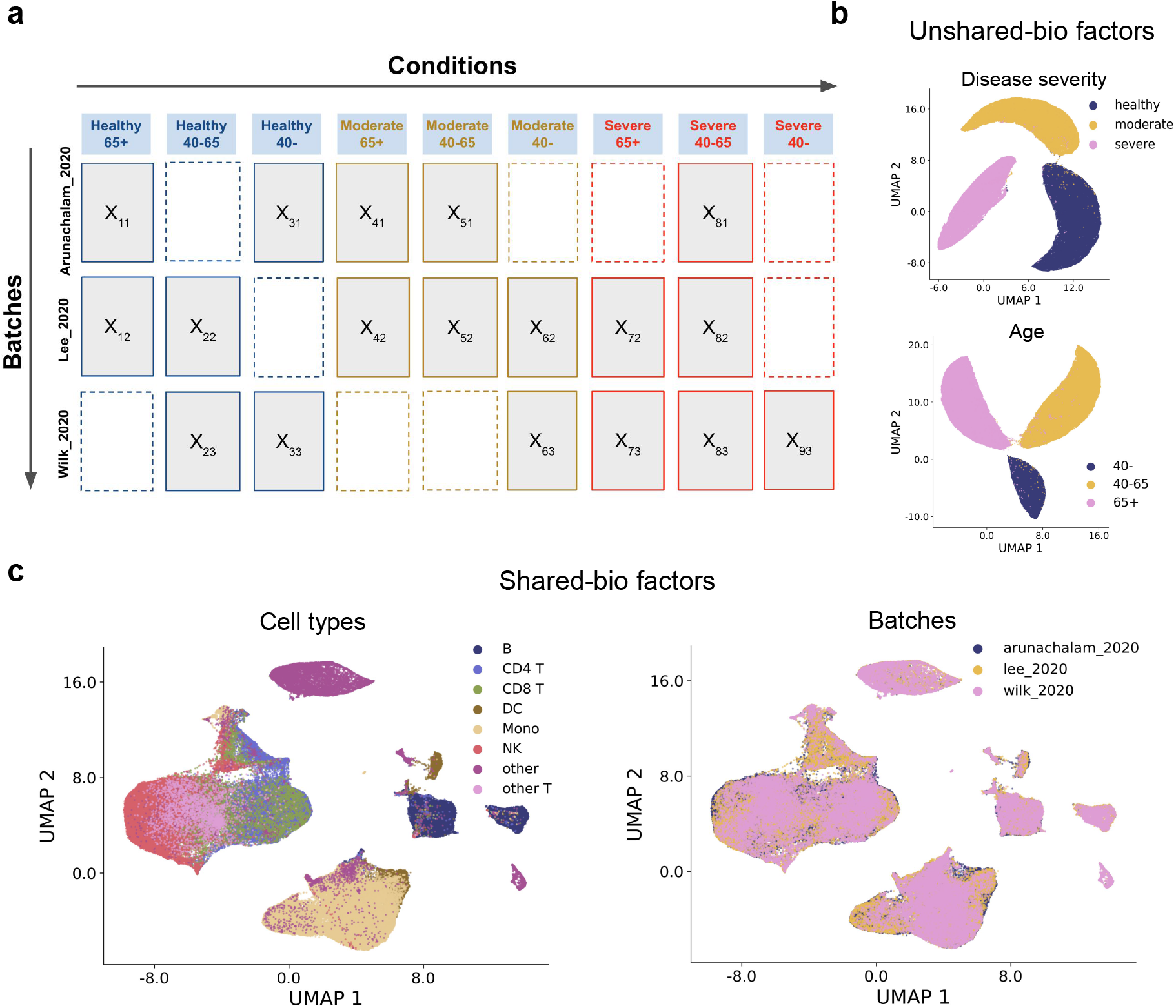
Test results on COVID-19 dataset. **a**. The arrangement of count matrices in the datasets, the matrices are grouped by conditions (columns) and batches (rows). Rectangles with dashed borders represent missing conditions for the corresponding batch. **b**. The UMAP visualization of unshared-bio factors, where the upper plot shows the factor that encodes disease severity condition, and the lower plot shows the factor that encodes age condition. Cells are colored according to their ground truth conditions. **c**. The UMAP visualization of shared-bio factors in scDisInFact, where cells are colored by (left) ground truth cell types and (right) batches.

We first visualize the gene expression data using UMAP before applying scDisInFact. From the visualization, we observed a strong batch effect among different studies (Figs. S7a,b). In addition, the distributions of cells under different conditions also show variation due to the condition effect (Figs. S7c,d). We then trained scDisInFact on the dataset, and visualized the shared and unshared biological factors using UMAP (Figs. 5b,c). The visualization shows that the shared-bio factors are aligned across batches and conditions while maintaining the same level of cell type separation as in individual batches (Fig. 5c, Fig. S7a), and the unshared-bio factors are also grouped according to their corresponding condition types (Fig. 5b).

We then tested the perturbation prediction of scDisInFact. Similar to the test on simulated datasets, we design tests separately for the prediction of condition combinations that are seen (*in-sample* test) and unseen (*out-of-sample* test) in the training set. In *in-sample* test, we trained scDisInFact on all available count matrices (Fig. S8a), whereas in *out-of-sample* test, we held out all count matrices under condition <*moderate*, *40-65*> (*X*_51_ and *X*_52_ in Fig. S8b) and train scDisInFact on the remaining count matrices. In both tests, given an input count matrix, we predict the gene expression data of the same cells under the condition <*moderate*, *40-65*> and batch "Lee_2020" (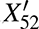 in Figs. S8a,b). Depending on whether condition effect and batch effect exist between input and predicted count matrices, we again categorize the prediction tests into 6 scenarios, similar to the test on simulated datasets. When the predicted and input count matrices are from the same batch, we test the prediction of condition effect of *disease severity* (arrows 1 in Figs. S8a,b), *age* (arrows 2 in Figs. S8a,b), and *disease severity & age* (arrows 3 in Figs. S8a,b). Similarly, when the predicted and input count matrices are from different batches, we also test the prediction of these three combinations of condition effects (arrows 4,5,6 in Fig. S8a,b).

We again compare scDisInFact’s perturbation prediction performance with scGen and scPreGAN. The detailed settings of the training, input, and predicted conditions for all methods are summarized in Table S7. We use the held-out count matrix corresponding to condition <*moderate*, *40-65*> and batch "Lee_2020" (*X*_52_ in Figs. S8a,b) as the gold-standard counts for scGen and scPreGAN and its denoised version for scDisInFact, and calculate cell-type-specific MSE, *R*^2^, and Pearson correlation scores between predicted and gold-standard counts (Methods), shown in Fig. 6. In both *in-sample* test and *out-of-sample* test, scDisInFact outperforms scGen and scPreGAN across all 6 prediction scenarios, where the improvement scDisInFact brings is more pronounced for the tasks that involve prediction of batch effects, compared to the tasks of predicting only condition effects. This can be attributed to scDisInFact’s ability to model both condition effects and batch effects, whereas scGen and scPreGAN model the condition effect without considering the batch effect.

**Figure 6.**
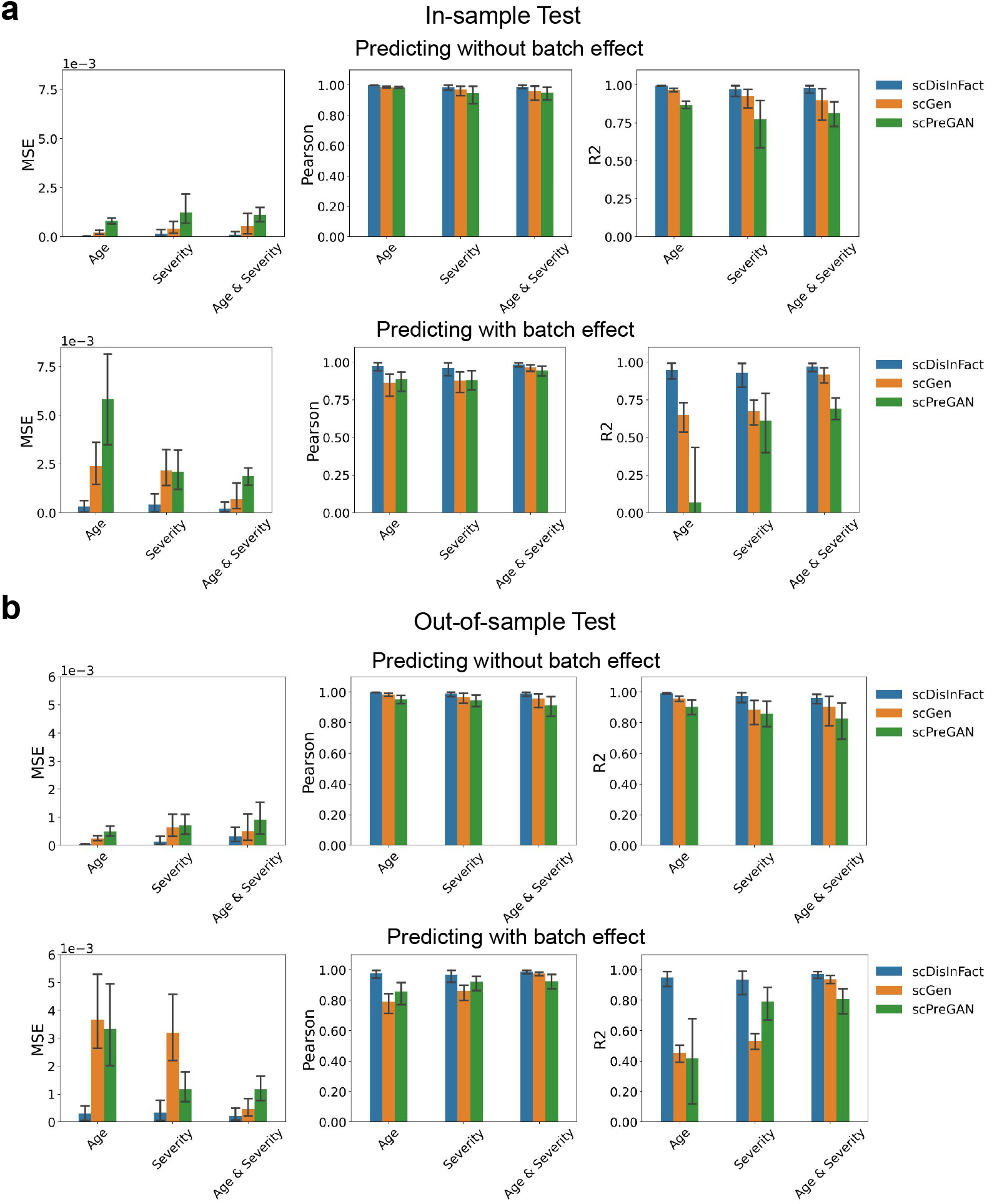
Perturbation prediction result on COVID-19 dataset. **a**. Barplots showing the perturbation prediction accuracy of scDisInFact and baseline methods in in-sample test. The upper row shows the result where the input and predict count matrices are from the same batch, and the lower row shows the result where the input and predict count matrices are from different batches. (b) Barplot showing the perturbation prediction accuracy of scDisInFact and baseline methods in out-of-sample test. Plots are organized following **a**.

## 3 Discussion

We presented scDisInFact, a deep learning framework that models multi-batch multi-condition scRNA-seq datasets. scDisInFact is a unified framework for three prominent tasks in disease study: (1) the disentanglement of biological factors and removal of batch effect, (2) detection of condition-associated key genes, and (3) prediction of gene expression data under conditions where no data is measured. The last two tasks are enabled by achieving the goal of the first task, the disentanglement of variations in a multi-batch multi-condition scRNA-seq dataset. While these tasks were conducted separately in existing work, they are related to each other and can benefit from a unified framework that outputs consistent results for each task. That scDisInFact performs a comprehensive disentanglement of various types of information in the dataset is key to its multi-task ability.

The extensive tests conducted on simulated and real datasets support that scDisInFact has superior performance than the baseline methods that can only conduct one task. Not only scDisInFact gains better performance on these tasks, but it is also more versatile than existing methods in each task which can lead to much broader applications under realistic scenarios. For batch effect removal, scDisInFact removes only batch effects and preserves biological differences across data matrices; For condition-associated key gene detection, not only scDisInFact can output CKGs at a high level, but the perturbation prediction results can also be used to find genes that are differentially expressed in specific cells from one condition combination to any other condition combination. For perturbation prediction, scDisInFact models multiple condition types associated with the donors and can predict data from a condition combination to any other combination under study. This enables applications in complex scenarios like predicting the effect of combinations of multiple drugs.

While scDisInFact performs well in the reported scenarios which are applicable to a wide range of real datasets, there are certain scenarios that can pose challenges to disentanglement methods. As shown in our results, when each batch of cells is measured under only one condition, there is not enough information to fully disentangle batch and condition effect. Measuring data from multiple conditions in the same batch can largely ameliorate this problem, and we recommend that wet-lab experimental design takes this into consideration.

## 4 Methods

### 4.1 Loss function and training procedure of scDisInFact

scDisInFact uses a combination of loss terms to accomplish the given tasks. Firstly, scDisInFact reconstructs the input gene expression data from the decoder by minimizing the evidence lower bound (ELBO) loss, following the design of variational autoencoder. We denote the input gene expression data, shared-bio factors, unshared-bio factors, and batch factors of each cell as **x**, **z***_s_*, **z***_u_*, and *b*, respectively. There can be multiple unshared-bio factors **z***_u_*s, each corresponding to one condition type (Fig. 1b). For clarity of explanation, we describe the case where only one condition type exists in the Method section, and the multiple-condition-type case can be easily generalized from one condition type. The ELBO loss for datasets with one condition type follows:

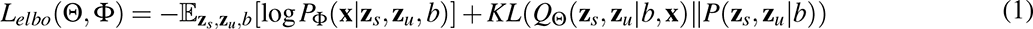

*Q*_Θ_(*·*) is the encoder network with parameters Θ that model the posterior distribution of **z***_s_* and **z***_u_* given **x** and *b*. *P*_Φ_(*·*) is the decoder network with parameter Φ that models the likelihood function of **x**. *P*(**z***_s_,* **z***_u_|b*) is the prior distribution of **z***_s_* and **z***_u_*. Since **z***_s_*, **z***_u_*, and *b* model factors that correspond to independent sources of variation, we can factorize *Q*_Θ_(**z***_s_,* **z***_u_|b,* **x**) into *Q*_Θ_(**z***_s_|***x***, b*)*Q*_Θ_(**z***_u_|***x***, b*), and *P*(**z***_s_,* **z***_u_|b*) into *P*(**z***_s_|b*)*P*(**z***_u_|b*) according to mean-field approximation^26^. We use a shared encoder to model *Q*_Θ_(**z***_s_|***x***, b*), and unshared encoders to model *Q*_Θ_(**z***_u_|***x***, b*). Then the ELBO loss can be simplified as

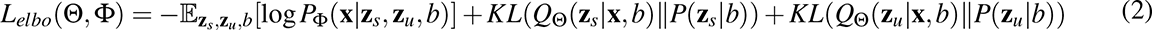

where *P*(**z***_u_|b*) and *P*(**z***_s_|b*) follow a standard normal distribution *N*(**0**, **I**). We model the likelihood function *P*_Φ_ (**x***|***z_s_**, **z_u_** *, b*) using a negative binomial distribution 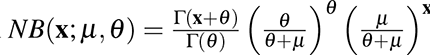, where *µ* and *θ* are the mean and dispersion parameters of the distribution^27,28^. The decoder learns the gene-specific *µ* and *θ* for each cell. By minimizing ELBO loss, scDisInFact is trained to extract the main biological variation in the dataset into latent factors and generate correct gene expression data from latent factors.

We train each unshared encoder along with its corresponding linear classifier to predict the condition of each cell, and calculate cross-entropy loss *L_ce_*(*y*_cond_*, y*_class_) using the classifier’s outputs *y*_class_ and condition labels *y*_cond_. By minimizing *L_ce_*(*y*_cond_*, y*_class_), the unshared encoder is guided to extract the condition-related biological variations from the input data. In addition to the cross-entropy loss, we also add a circle contrastive loss^29^ (*L_contr_*(**z***_u_, y*_class_)) on the output of the unshared encoder **z***_u_*. The circle contrastive loss also aims to separate cells from different conditions (Supplementary Note 1) which further improves the classification accuracy of the model^30^.

To remove the batch effect from the shared-bio and unshared-bio factors, we apply maximum mean discrepancy (MMD) loss on **z***_s_* and **z***_u_*. The MMD loss calculates the degree of mismatch between two distributions, and was used to align the latent embedding of cells across batches^11,31^ (Supplementary Note 2). **z***_s_* is irrelevant to conditions and batches, which means that **z***_s_* from different batches and conditions should follow the same distribution. We added MMD loss on **z***_s_* to align its distribution across batches and conditions. We denote the set of batches under condition label *c* as *B_c_*, and the total number of condition labels as *C*. The MMD loss on **z***_s_* is the sum of MMD losses between **z***_s_* from a reference batch and condition 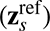 and **z***_s_* from each of the remaining batches and conditions 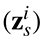,

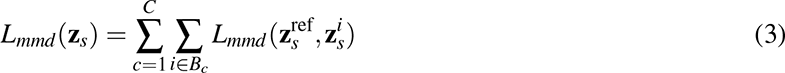

We used the **z***_s_* of the 1st batch and condition label as 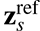 in all our tests. The unshared-bio factors encode the condition-related biological variations, which means **z***_u_* from batches of the same condition label follow the same distribution, and **z***_u_* from batches of different condition labels follow different distributions. For each condition label *c*, we select **z_u_** from one reference batch 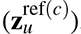 and calculated one MMD loss between **z_u_** from the reference batch and the other batches. The final MMD loss for the unshared-bio factors is the sum of MMD loss across condition labels, following

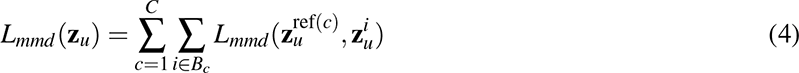

We used the 1st batch of each condition label *c* as the reference batch ref(c) in all our tests.

To enforce the disentanglement of shared-bio and unshared-bio factors, we apply total correlation loss^21,32^ on **z***_s_* and **z***_u_*, which was used to enforce independence among latent variables. If **z***_u_* and **z***_s_* are independent (disentangled), we have *Q*_Θ_(**z***_u_,* **z***_s_*) = *Q*_Θ_(**z***_u_*)*Q*_Θ_(**z***_s_*). The total correlation loss measures the KL-divergence between *Q*_Θ_(**z***_u_,* **z***_s_*) and *Q*_Θ_(**z***_u_*)*Q*_Θ_(**z***_s_*)^32^. Following Factor-VAE^21^, we minimize the KL-divergence using a discriminator network, and transform the problem into a two-player minimax game between encoder and discriminator, similar to GAN^33^. We first create two sample sets: 1. we directly concatenate **z***_s_* and **z***_u_* from each mini-batch as the sample set from *Q*_Θ_(**z***_u_,* **z***_s_*); 2. we shuffle the **z***_u_* in the mini-batch before concatenating it with **z***_s_* to create sample set from *Q*_Θ_(**z***_u_*)*Q*_Θ_(**z***_s_*). We then train the discriminator network to predict the origin of the samples (the first or the second set), and train the encoder to fool the discriminator network. Denoting *D* as the discriminator network, the objective function follows

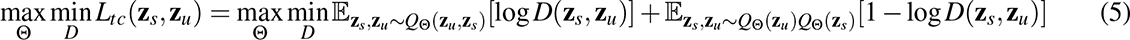

We transform the first layer of each unshared encoder into a feature (gene) selection layer through group lasso loss^34,35^. We represent the weight matrix of the first layer by **W** = [**w**_1_, **w**_2_*, · · ·,* **w***_d_*], where *d* is the number of input dimensions (genes), and **w***_i_* is the *i*th column vector of **W** connecting to the *i*th gene. The group lasso loss 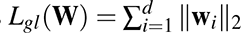 penalizes the number of non-zero **w***_i_*s, thus the first layer of each unshared encoder is forced to select the most discriminative genes as the condition-associated key genes (CKGs) of the corresponding condition type.

The objective function of scDisInFact consists of a weighted combination of losses above, which follows

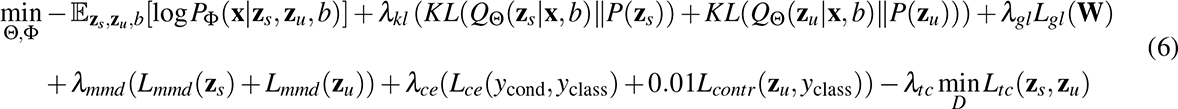

where *λ_kl_*, *λ_ce_*, *λ_gl_*, and *λ_tc_* are the weights of the losses.

### 4.2 Training algorithm

We update the model parameter using in an alternating manner using stochastic gradient descent. For each iteration, the parameter update of scDisInFact is separated into 3 steps. We first fix the parameters of the unshared encoder, classifier, and discriminator networks, and update the parameters of the shared encoder and decoder networks through stochastic gradient descent. The loss function (Equ. 6) is then simplified into:

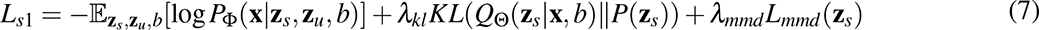

Then we fix the parameters of the shared encoder and discriminator networks, and update the parameters of the unshared encoder, classifier, and decoder networks through stochastic gradient descent. The loss function (Equ. 6) is then simplified into:

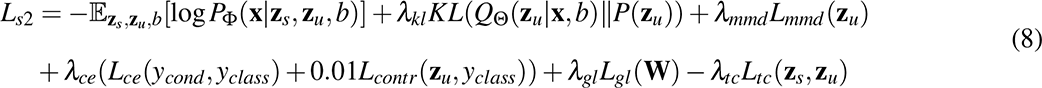

Finally, we fix the parameters of all networks except the discriminator network, and update the parameters of the discriminator network through stochastic gradient descent. The loss function (Equ. 6) is then simplified into:

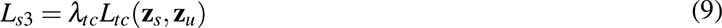

The algorithm iterate until the objective function (Equ. 6) converges.

### 4.3 Condition-associated key gene (CKG) detection

The weight matrix **W** in the first layer of each unshared encoder is used to extract the CKGs of its corresponding condition type. Each column vector of **W** is connected to one input gene, and we used the *ℓ*_2_-norm of each column vector as the score of the corresponding gene. For gene *i*, the corresponding score *s_i_* is calculated as *s_i_* = ∥**w***_i_*∥_2_, where **w***_i_* is the *i*th column vector of **W**. A higher *s_i_* score means that gene *i* is more likely to be a CKG.

### 4.4 Prediction of condition effect on gene expression data

Given input gene expression data under one condition, scDisInFact is able to predict the corresponding data under other input conditions. We illustrate the prediction procedure of scDisInFact using the example in Fig. S1b. Fig. S1b describes a case where the dataset has 3 condition labels (condition 1, 2, and 3 in unshared-bio factors) and 6 batches (6 dimensions in batch factors). In the example, scDisInFact takes as input a count matrix under condition 1 and batch 6, and predicts the count under condition 2 and batch 1 (Fig. S1b). To do the prediction, we need to calculate new unshared-bio factors through latent space arithmetics^17^(Fig. S1b (left)). The latent space arithmetics includes two steps: (1) we calculate the shifting vector *δ* that measures the difference between the mean unshared-bio factors under *condition 1* 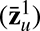 and *condition 2* 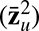, following 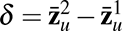; (2) For all cells under *condition 1*, we shift their unshared-bio factors 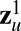 by *δ*, following 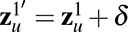. Then we need to update the batch factor to match the predicted batch. In Fig. S1b, since the predicted count belongs to batch 1, we assign 1 to the 1st dimension of the batch factor and 0s to the remaining dimensions. We keep the shared-bio factors to be the same as the shared-bio factors do not encode any condition or batch effect. Finally, we feed the original shared-bio factors, the updated unshared-bio factors, and the updated batch factors into the decoder, and use the decoder output *µ* as the predicted counts. Decoder model the gene expression data using a negative binomial distribution, and the output *µ* is interpreted as the denoised gene expression data.

### 4.5 Hyper-parameter selection

The main hyper-parameters of scDisInFact include the latent dimensions of shared-bio and unshared-bio factors, and the weight parameters in the loss function. The most common way of hyper-parameter selection on such a model is conducting a grid search of the hyper-parameter on the held-out dataset. Given a dataset, one can separate 10 percent of cells from each batch to create a testing dataset, train the model on the remaining dataset, and check the losses on the testing dataset (e.g. log-likelihood loss, classification loss, etc). However, the grid search is extremely time-consuming given a large set of hyper-parameters or a large input dataset. Here we also provide a recommended hyper-parameter setting, which was obtained from extensive tests on both real and simulated datasets. We recommend the latent dimensions of the shared encoder to be 8 and of unshared encoders to be 2 *∼* 4. The recommended weights are *λ_kl_* = 10*^−^*^5^, *λ_mmd_* = 10*^−^*^4^, *λ_ce_* = 1, *λ_gl_* = 1, and *λ_tc_* = 0.5. We used the recommended hyper-parameters for all our test results in the manuscript. Users can manually tune the hyper-parameters around the recommended setting instead of conducting a comprehensive grid search, and it should provide a reasonable result on most of the dataset.

We also provide the detailed parameter of neural networks in scDisInFact. The shared encoder is a 3-layer fully connected neural network. The first 2 layers have the same structure, where each layer consists of a linear layer followed by a ReLu activation function and a dropout layer (“linear”-“ReLu”-“dropout”, 128 output neurons for each layer). The last layer has two linear networks that produce the mean and variance of the shared-bio factors separately. The unshared encoder has 2 layers. The first layer also follows the “linear”-“ReLu”-“dropout” structure (128 output neurons), and the last layer has two separate linear networks for the mean and variance of the unshared-bio factors. The decoder is a 3-layer fully connected neural network. The first 2 layers also follow the “linear”-“ReLu”-“Dropout” structure (128 output neurons for each layer). The last layer includes two linear networks that separately produce the mean and dispersion parameters of the data distribution. The dropout rate of all dropout layers in scDisInFact is 0.2.

### 4.6 Simulation procedure

We simulated multi-batch scRNA-seq datasets using SymSim^36^, and then added condition effect on the simulated dataset. We selected *m^c^* genes as the CKGs for each condition type *c* (CKGs of different condition types have no overlap), and added the condition effect of condition type *c* on the corresponding CKGs in the Symsim-generated data. We denote the Symsim-generated count matrix as **X***_obs_*, where **X***_obs_*[*i, g*] corresponds to the expression level of a CKG *g* in cell *i*. Then the condition effect was added as a uniform distribution on **X***_obs_*[*i, g*] with lower bound *ε −* 1 and upper bound 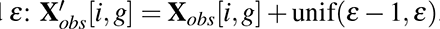. *ε* is the *perturbation parameter* that controls the strength of the condition effect. For each condition type, multiple conditions can be generated with different condition labels. For example, one can generate count matrices under the control and stimulated conditions, where no condition effect is added on the control condition, and the condition effect on the count matrices in the stimulated condition is added following 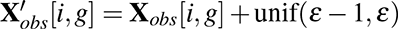. The modeling of batch effect is already included in Symsim simulator^36^.

We generated 9 simulated datasets with 2 condition types and 2 data batches (8 count matrices for each dataset, Fig. 2a). The 2 condition types respectively have condition labels (1) control and stimulation, (2) healthy and severe. We set the first 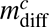 genes to be the CKGs of conditions control and stimulation. We set the 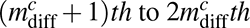 genes to be CKGs of conditions healthy and severe. We first generate 2 batches of count matrices using Symsim. Then we evenly separate cells of each batch and the corresponding gene expression data into 4 conditions: (control, healthy), (control, severe), (stimulation, healthy), and (stimulation, severe). For each chunk of the gene expression data, we add condition effect according to its condition group. We do not add condition effect for the gene expression data under (control, healthy), whereas we add condition effect to the corresponding CKGs for gene expression data under either stimulated or severe conditions. The 9 datasets are generated with different simulation parameters to model different strengths of condition effect. We used 3 CKG numbers 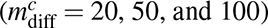, and generated 3 datasets under each CKG number with 3 perturbation parameters (*ε* = 2, 4 and 8). The detailed simulation parameter settings of 9 datasets are included in Table S1.

### 4.7 Pre-processing steps of real datasets

scDisInFact can be trained directly on the raw scRNA-seq dataset, no pre-processing step is required before running scDisInFact. However, directly training on raw scRNA-seq data can be time-consuming as the feature dimension (number of genes) in the raw data can be extremely high (20,000*∼*30,000). Some additional gene filtering steps that remove genes with low expression levels are also recommended, as they can greatly improve the running speed of the model.

#### 4.7.1 Pre-processing steps of GBM dataset

In the GBM dataset, the authors obtained multi-batch scRNA-seq data from the GBM resections of patients with different drug treatments. We selected 21 count matrices from 6 GBM patient batches with and without panobinostat drug treatment (respectively named *0.2 uM panobinostat* and *vehicle (DMSO)*), where 16 matrices were under *vehicle (DMSO)* condition and 5 matrices were under *0.2 uM panobinostat* condition (see Table S5 for detailed information on the selected batches). We remove the genes that have counts in less than 100 cells across all batches, and obtained count matrices with 74777 cells and 19949 genes. The original paper annotated the cells into tumor and non-tumor cells, which is a very high-level annotation. We further annotated the non-tumor cells into Myeloid, Oligodendrocytes, and Other cells using their marker genes^1^ for each batch separately.

#### 4.7.2 Pre-processing steps of COVID-19 dataset

We selected COVID-19 scRNA-seq studies from the recent summary paper^16^ (data downloaded from https://atlas.fredhutch.org/fredhutch/covid/), and used the count matrices stored in “arunachalam_2020_processed.HDF5”, “wilk_2020_processed.HDF5”, and “lee_2020_processed.HDF5”. We followed its standard of disease severity classification, and select the data under the condition “healthy”, “moderate”, and “severe”. We categorized the ages of patients in the studies into groups of “40-” (below 40), “40-65” (between 40 and 45), and “65+” (above 65). Then, we selected the genes that are shared among all three studies. We did not conduct further filtering steps of genes and cells in these studies.

### 4.8 Running details of baseline methods

#### 4.8.1 Running details of scINSIGHT

When running scINSIGHT on simulated datasets, we preprocessed the dataset following the Seurat normalization steps (as was recommended in scINSIGHT online tutorial: https://github.com/Vivianstats/scINSIGHT/wiki/scINSIGHT-vignette) and did not conduct gene filtering steps on the datasets. The main hyper-parameters of scINSIGHT include the number of common gene modules (*K*) and the number of condition-specific gene modules (*K_j_*). We used the default setting of *K*, where scINSIGHT searched through *K* = 5, 7, 9, 11, 13, 15 and selected the best performing *K*.

When using scINSIGHT to learn shared-bio and unshared-bio factors, we used the default setting of *K_j_* (*K_j_* = 2) and remaining hyper-parameters. scINSIGHT learns a factorized common module and a factorized condition-specific module. The common module includes a membership matrix **V** and expression levels matrices **W***_b_*_2_ (*b* = 1, 2*, · · ·, B* for *B* batches). **W***_b_*_2_ is equivalent to the shared-bio factors of cells under batch *b*. The condition-specific module includes membership matrices **H***_c_* (*c* = 1, 2*, · · ·,C* for *C* conditions) and expression levels matrices **W***_b_*_1_ (*b* = 1, 2*, · · ·, B* for *B* batches). The unshared-bio factors are encoded in both **W***_b_*_1_ and **H***_c_*. We calculated the dot product of **W***_b_*_1_ and **H***_c_*, and obtained cell-by-gene matrices 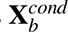 that only encoded the condition-related information. We treated 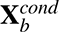 as the unshared-bio factors of scINSIGHT.

When using scINSIGHT to detect CKGs, we set *K_j_* = 1 such that the membership matrices **H***_c_* shrink into 1D vectors with length equal to the number of genes. We then treated **H***_c_*s as the importance score of genes under condition *c*. The genes that have important scores varing the most across conditions should be the CKGs. We calculate the variance of gene *i* under different conditions following

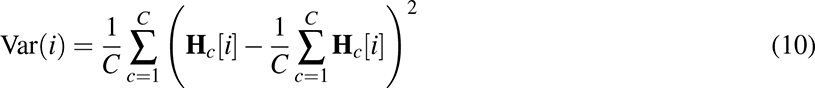

We then normalize the variances of genes to the range of 0 and 1, which are used as the scoring of CKGs from scINSIGHT.

#### 4.8.2 Running details of scGen and scPreGEN

We ran scGen following the steps and parameter setting in its online tutorial (https://scgen.readthedocs.io/en/stable/tutorials/scgen_perturbation_prediction.html). We ran scPreGAN following the parameter setting in its online repository (https://github.com/XiajieWei/scPreGAN-reproducibility).

In perturbation prediction tests, we trained scGen and scPreGAN on the datasets that are only normalized with library size. As the test requires the model to generate the count matrix as close as the gold-standard count matrix, and additional preprocessing steps introduce unnecessary bias. Both scGen and scPreGAN are designed for the perturbation prediction task where only one condition type is involved. To predict the condition effect on datasets where two condition types are involved (simulated and COVID-19 datasets), we ran both methods on a subset of count matrices where one condition type is fixed and another condition varies, and the methods are trained to predict the condition effect of the changed condition. To further predict the joint effect of two condition types (e.g. *disease severity* and *age* in COVID-19 dataset), two cascade models are needed for both methods, where the first model learns one condition effect and the second model learns the other condition effect. For the test datasets, the detailed settings of training data used in both methods are described in Tables S2,3,6-8.

### 4.9 Evaluation metrics and gold-standard data

We evaluate the disentanglement of biological factors and technical batch effect using the ARI (adjusted rand index) and ASW-batch (batch-mixing average silhouette width) scores^37^. The ARI score measures the matching of latent space cluster result and ground truth cell type label. An ARI score ranges from 0 to 1, and a higher score means better separation of cell types in the latent space. The ASW-batch scores measure how well the cells of the same cell type are aligned among batches in the latent space^38^. The scores range between 0 and 1, and a higher score means a better alignment of batches and removal of batch effect.

To evaluate the CKGs detection accuracy, we use AUPRC (area under the precision-recall curve) score, where a higher AUPRC score means a better detection accuracy. The AUPRC is calculated using ground truth CKGs and the inferred CKG scores of each method.

When evaluating perturbation prediction, we use the denoised count matrix as gold-standard for scDisInFact. This is because the known matrices have technical noise given the procedure of data simulation, while the scDisInFact prediction is already denoised, thanks to the use of the negative binomial distribution in the likelihood function. Directly comparing scDisInFact prediction with the simulated counts would also induce the error caused by technical noise. We instead pass the simulated counts through the scDisInFact model to generate the denoised output, and compare the predicted counts with the denoised simulated counts instead. We still compare the prediction result of scGen and scPreGAN with the simulated count directly, since these methods did not model the technical noise in their prediction.

We evaluate the prediction accuracy of gene expression data using MSE (mean square error), Pearson correlation, and *R*^2^ scores. We calculate the scores on count matrices that are normalized against library size. MSE measures the difference between true and predicted count matrices on all input genes, where smaller values are better. Pearson correlation also measures the alignment of true and predicted count matrices. It ranges between -1 and 1, and a higher Pearson correlation means a better prediction result. *R*^2^ measures the coefficient of determination. A higher *R*^2^ score means a better matching between the predicted and true counts, and the maximum *R*^2^ score is 1. Since the gold-standard counts and the predicted counts are not from the same cell, there is no direct way to calculate MSE, Pearson correlation, and *R*^2^ for each cell. We instead calculate the score in a cell-type-specific manner. We calculate the centroid of each cell type using the mean gene expression data, and then measure the MSE, Pearson correlation, and *R*^2^ between the centroid gene expression value of predicted and gold-standard counts for each cell type.

## 5 Data availability

The datasets used in this study are all publicly available. The GBM dataset is accessible with the accession code GSE148842. The COVID-19 dataset is downloaded from the website with the link: https://atlas.fredhutch.org/fredhutch/covid/. The original data are also accessible with the accession number: GSE155673, GSE149689, and GSE150728.

## 6 Code availability

The code of scDisInFact is available on GitHub with the link: https://github.com/ZhangLabGT/scDisInFact.

## 7 Acknowledgements

This work was supported in part by the US National Science Foundation DBI-2019771 and National Institutes of Health grant R35GM143070. The authors would like to thank Dr. Yu Li for helpful discussions on this project.

## Supplementary Information

### Supplementary Figures and tables

**Figure S1.**
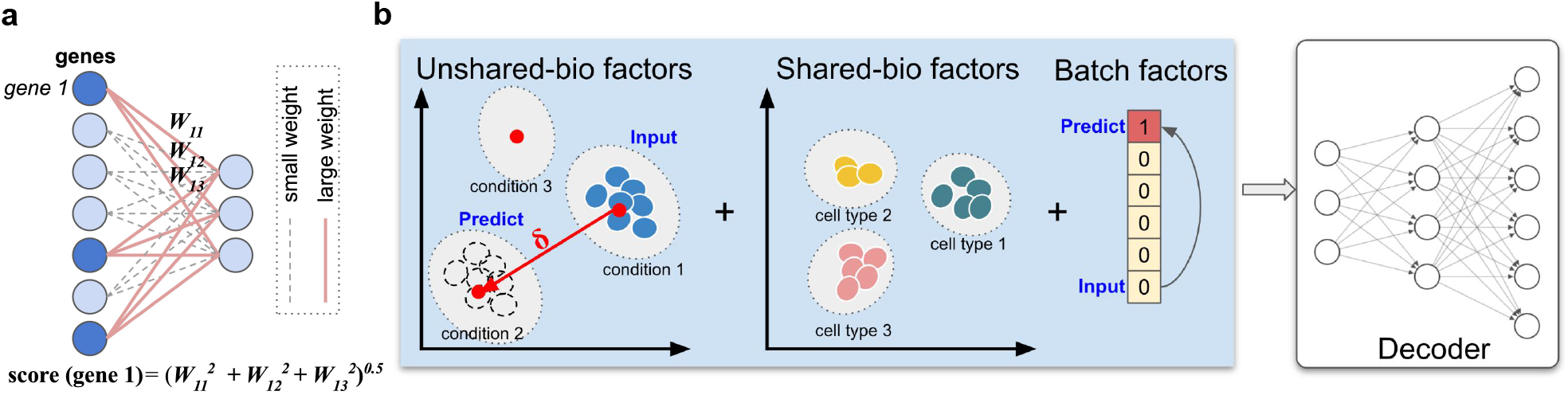
Key gene detection and perturbation prediction using scDisInFact. **a**. The first layer of the unshared encoder is designed to extract CKGs. **b**. By updating the unshared-bio factors and batch factors, scDisInFact allows for the perturbation prediction across conditions and batches.

**Figure S2.**
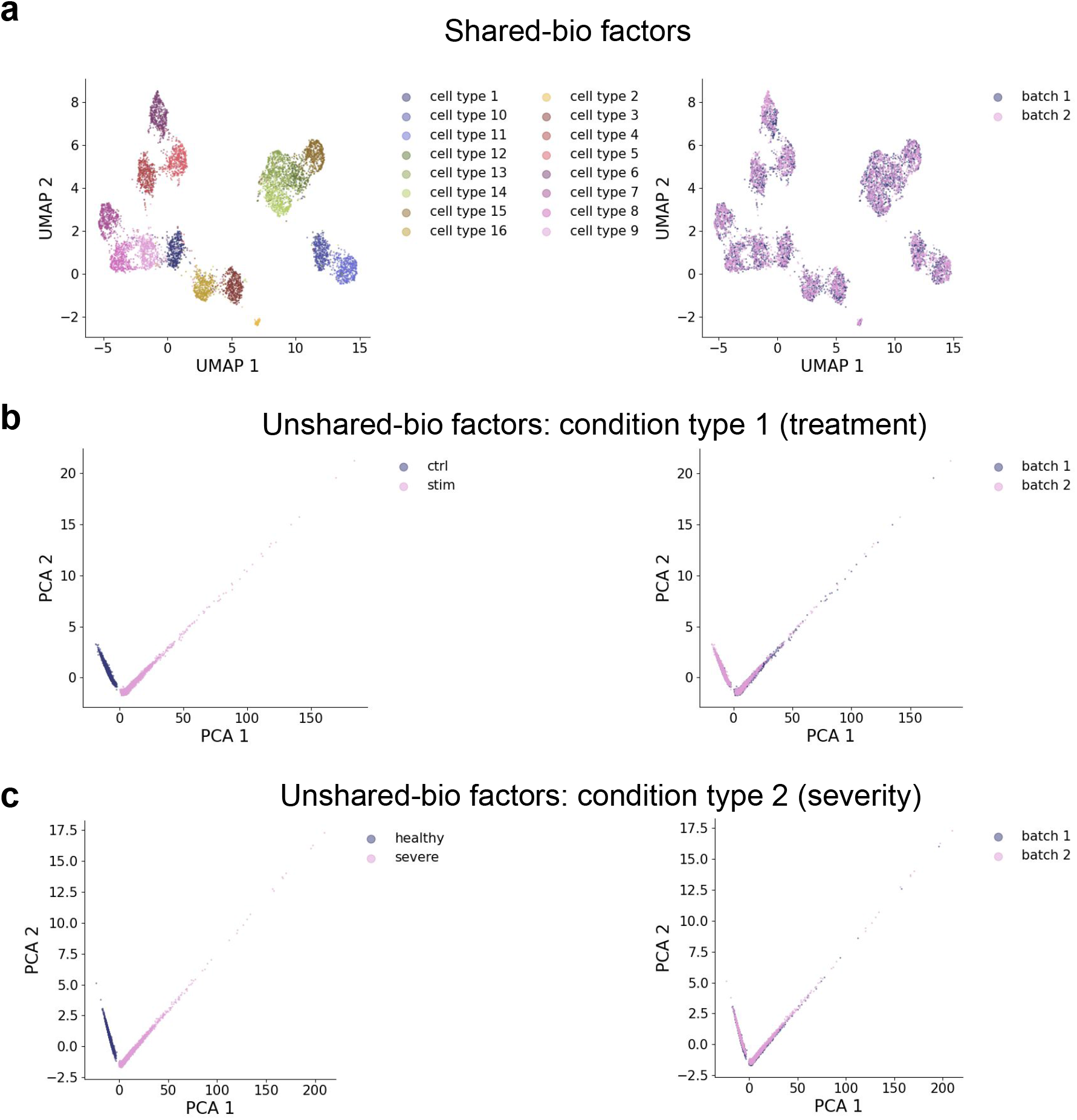
Latent factors of scDisInFact on one simulated dataset. **a**. The UMAP visualization of shared-bio factors, where cells are colored by (left) cell type identity and (right) batches. **b**. The PCA visualization of unshared-bio factors corresponds to the first condition type, where cells are colored by (left) conditions and (right) batches. **c**. The PCA visualization of unshared-bio factors corresponds to the second condition type, where cells are colored by (left) conditions and (right) batches.

**Figure S3.**
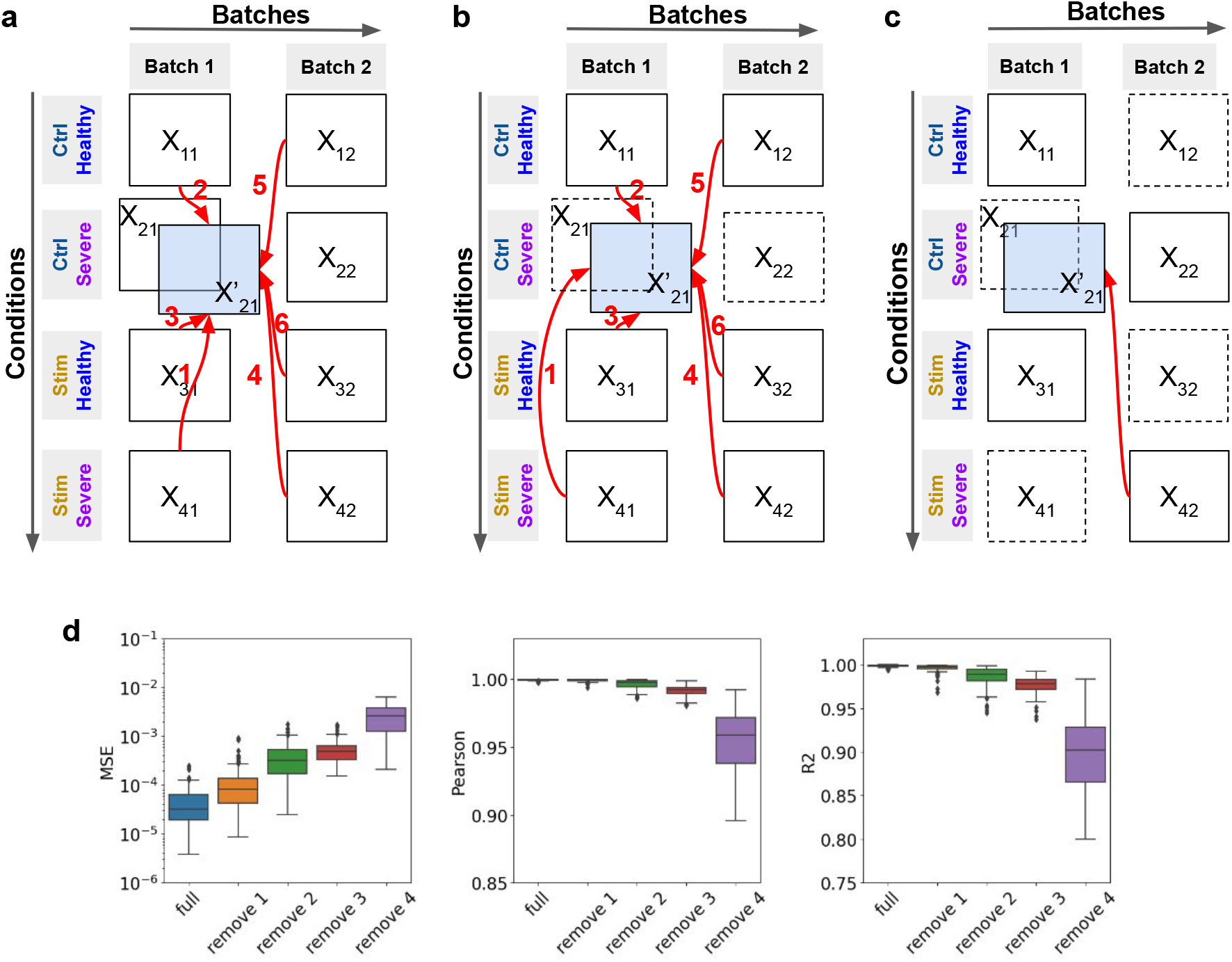
Additional perturbation prediction results on simulated datasets. **a**. Graphical illustration of in-sample test, where the red arrows represent 6 prediction categories. **b**. Graphical illustration of out-of-sample test, where the red arrows represent 6 prediction categories. **c**. Graphical illustration of prediction with different numbers of held-out matrices. Matrices with dashed borders are the held-out matrices when all 4 matrices are removed. **d**. prediction accuracy of scDisInFact with different numbers of held-out matrices, accuracy is measured with cluster-specific MSE, Pearson correlation, and *R*^2^ score.

**Figure S4.**
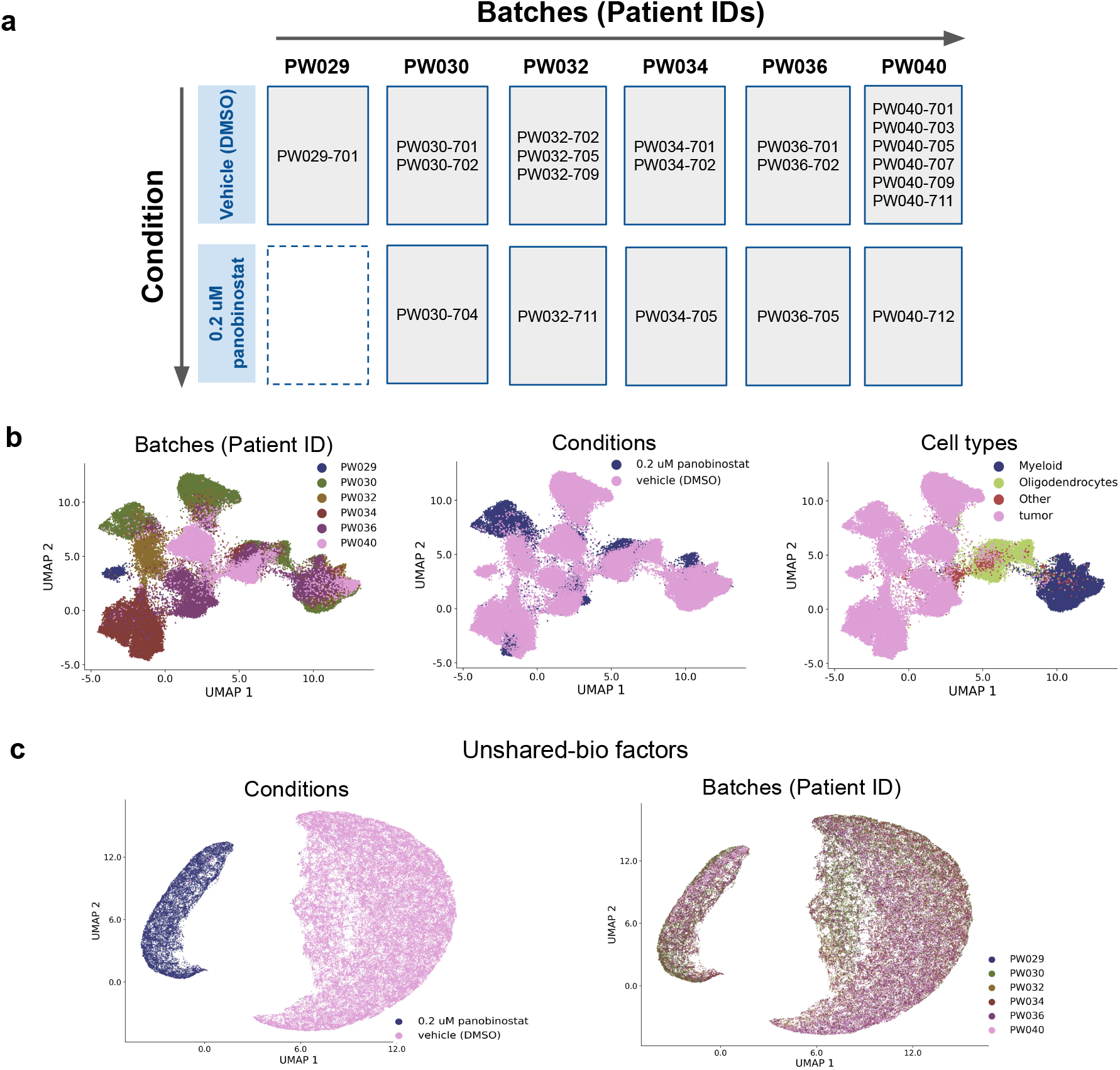
Additional test results on GBM dataset. **a**. The arrangement of count matrices in the dataset, the matrices are grouped by conditions (rows) and batches (columns). **b**. The UMAP visualization on the count matrices, where cells are annotated by (left) batch IDs, (middle) conditions, and (right) cell types. Cell-type labels are obtained from the original data paper. **c**. The UMAP visualization of the unshared-bio factors, where cells are annotated by conditions (left) and batches (right).

**Figure S5.**
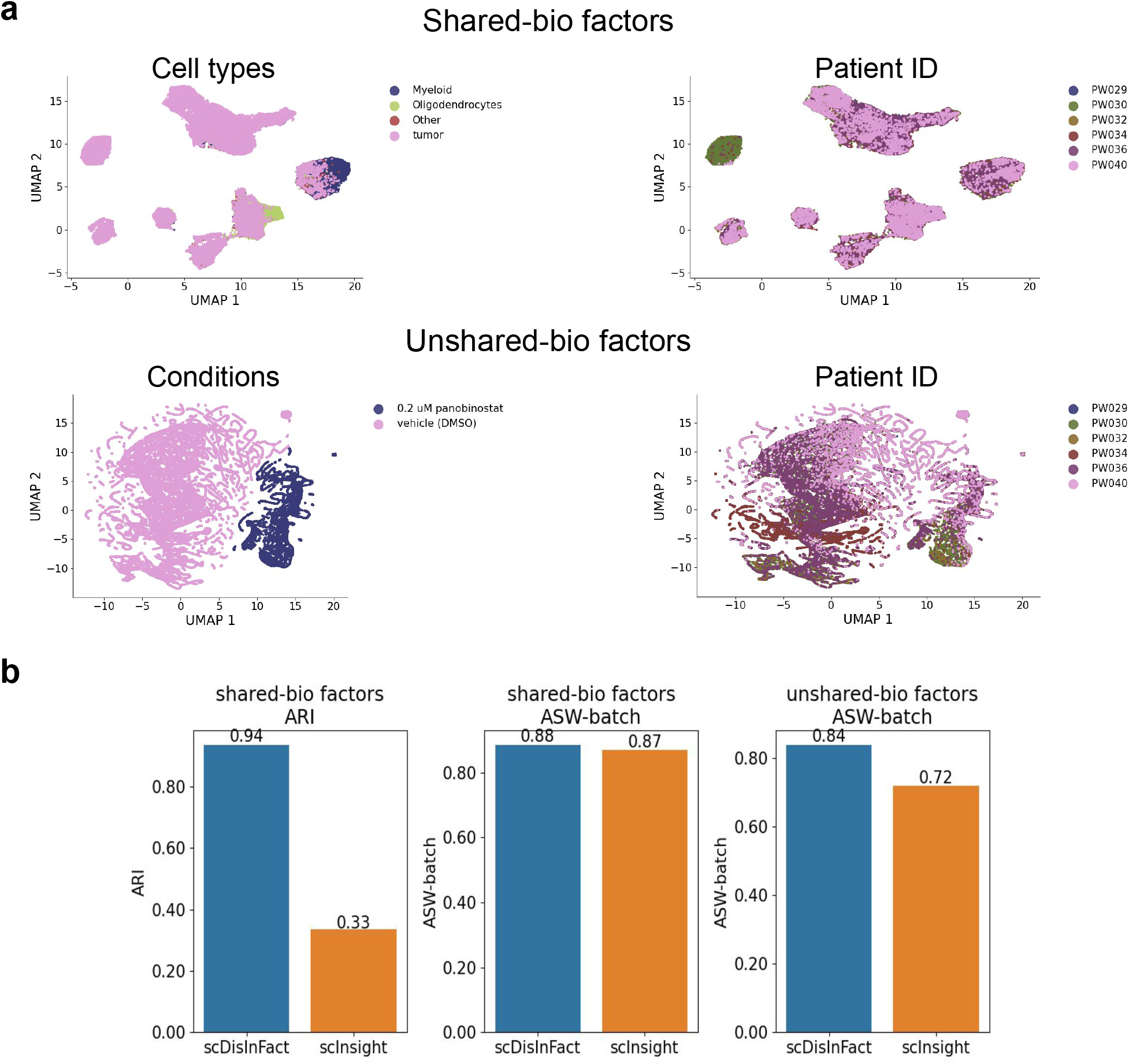
Additional test results on GBM dataset. **a**. The visualization of shared-bio and unshared-bio factors of scINSIGHT. **b**. The disentanglement scores of scDisInFact and scINSIGHT, which include the ARI (left) and ASW-batch (middle) scores for shared-bio factors, and the ASW-batch score (right) for unshared-bio factors.

**Figure S6.**
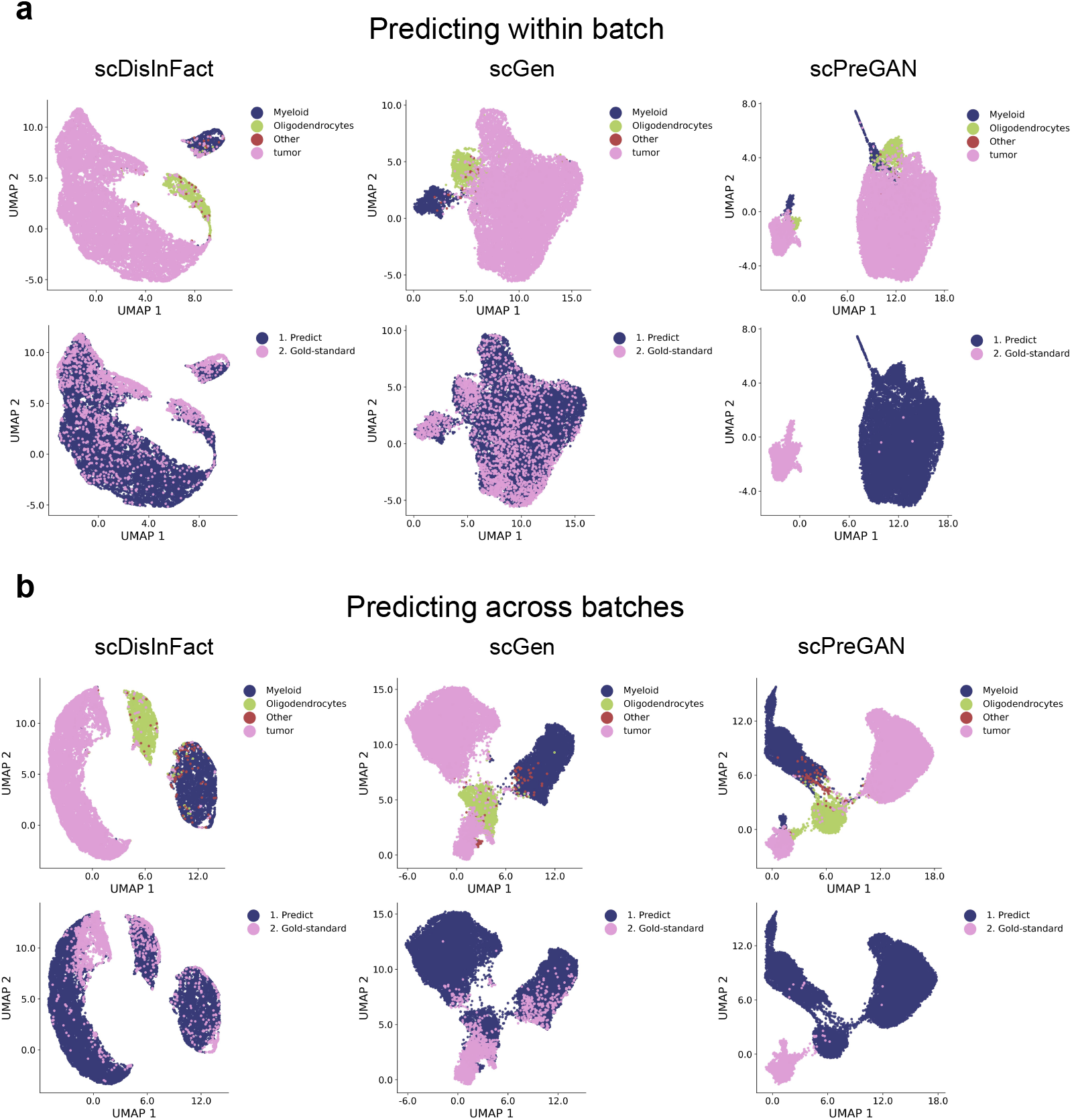
UMAP visualization of predicted and gold-standard matrices in the perturbation prediction test of GBM dataset. **a,b**. The UMAP visualization of the predicted and gold-standard matrices of scDisInFact, scGen and scPreGAN in (a) prediction task 1 (within batch) and (b) prediction task 2 (across batches). Cells are colored by (upper row) the original cell type and (lower row) the sources (predicted or gold-standard).

**Figure S7.**
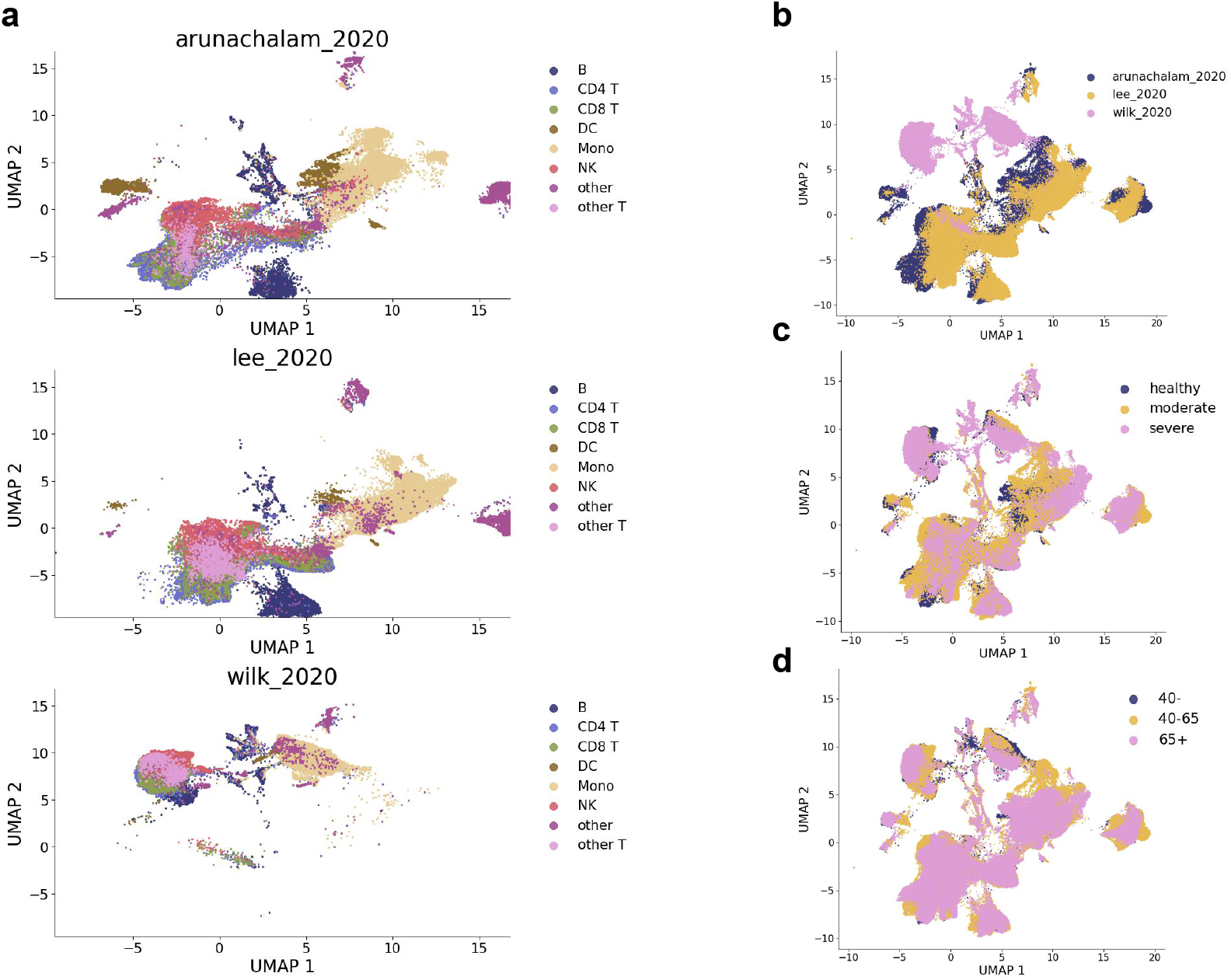
UMAP visualization of the count matrices in COVID-19 dataset. **a**. UMAP visualization of cells in different batches, where cells are colored by the input cell types. UMAP was calculated for cells from all three studies together to obtain the (x,y) coordinates for the cells in the UMAP space, but cells from different studies are shown in separate plots. **b-d**. UMAP visualization of cells, where cells are colored by (b) data batches, (c) disease severity, and (d) age groups.

**Figure S8.**
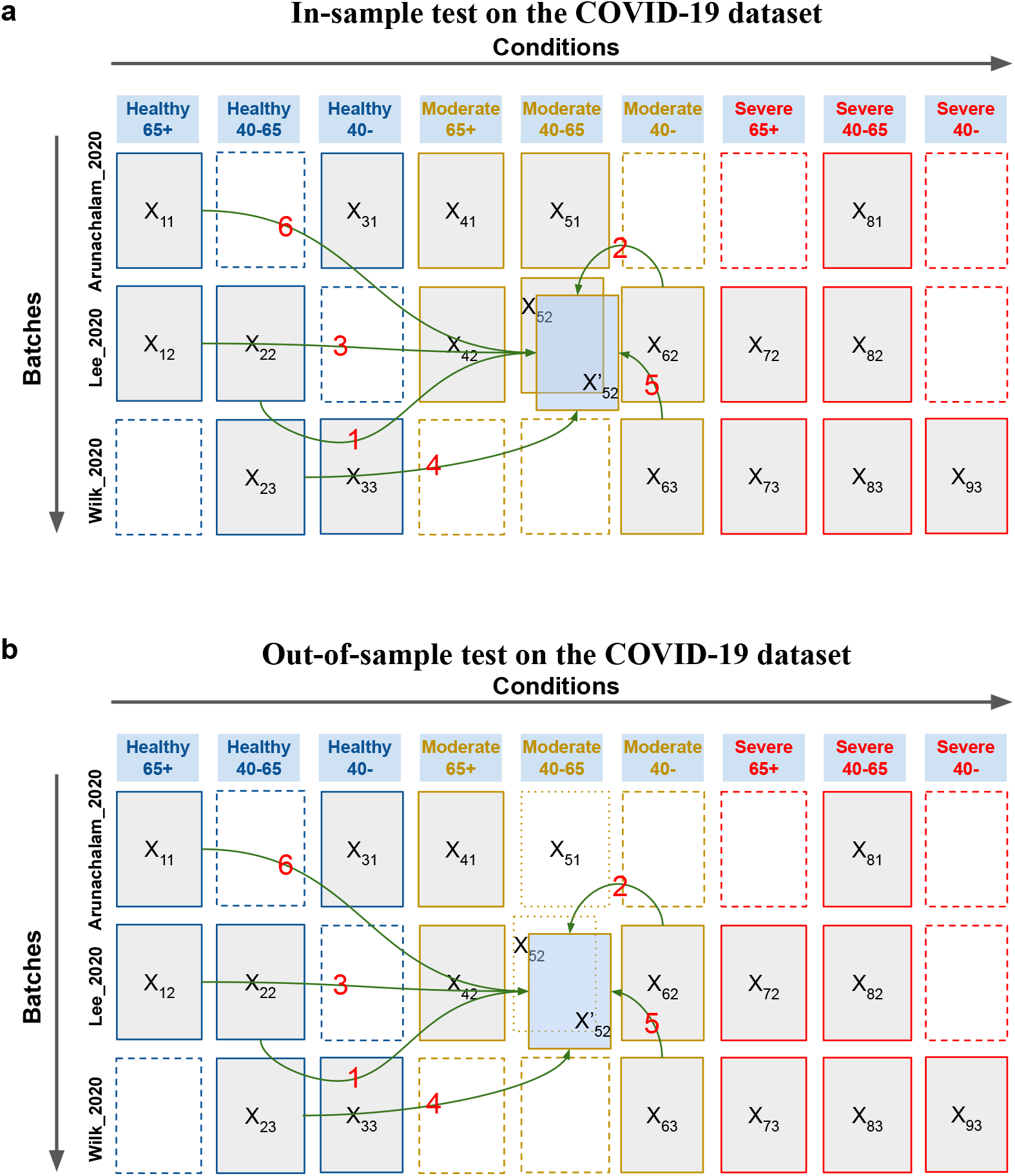
Graphical illustration of perturbation prediction test on COVID-19 dataset. **a**. The in-sample perturbation prediction test on COVID-19 dataset. **b**. The out-of-sample perturbation prediction test on COVID-19 dataset.

**Table S1.** The setting of simulation parameters in simulated datasets.

|  | Number of CKGs | Perturbation parameters |
| --- | --- | --- |
| Simulation 1 | 20 | 2 |
| Simulation 2 | 20 | 4 |
| Simulation 3 | 20 | 8 |
| Simulation 4 | 50 | 2 |
| Simulation 5 | 50 | 4 |
| Simulation 6 | 50 | 8 |
| Simulation 7 | 100 | 2 |
| Simulation 8 | 100 | 4 |
| Simulation 9 | 100 | 8 |

**Table S2.**
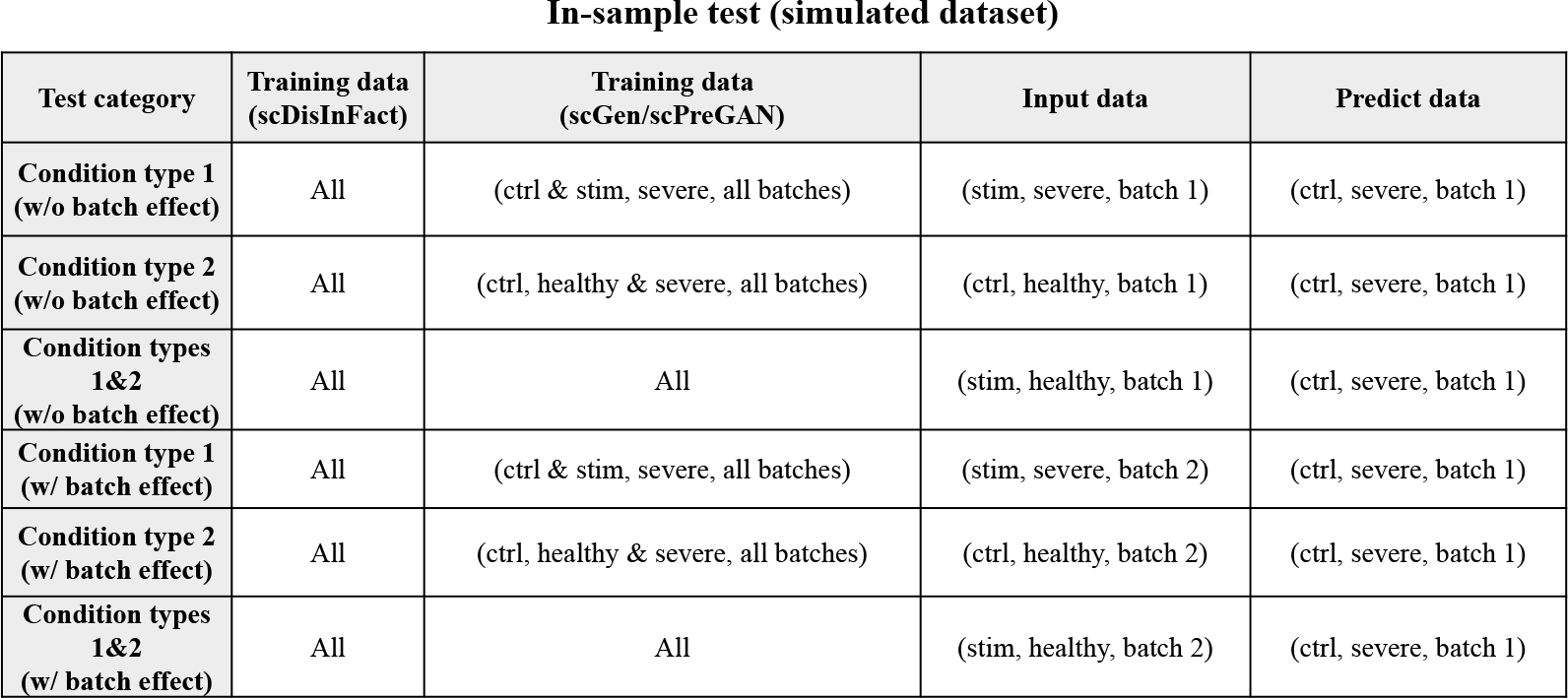
The setting of training data, input data, and predicted data of scDisInFact and baseline methods in *in-sample* prediction test on simulated datasets.

| Test category | Training data (scDisInFact) | Training data (scGen/scPreGAN) | Input data | Predict data |
| --- | --- | --- | --- | --- |
| Condition type 1 (w/o batch effect) | All | (ctrl & stim, severe, all batches) | (stim, severe, batch 1) | (ctrl, severe, batch 1) |
| Condition type 2 (w/o batch effect) | All | (ctrl, healthy & severe, all batches) | (ctrl, healthy, batch 1) | (ctrl, severe, batch 1) |
| Condition types 1&2 (w/o batch effect) | All | All | (stim, healthy, batch 1) | (ctrl, severe, batch 1) |
| Condition type 1 (w/ batch effect) | All | (ctrl & stim, severe, all batches) | (stim, severe, batch 2) | (ctrl, severe, batch 1) |
| Condition type 2 (w/ batch effect) | All | (ctrl, healthy & severe, all batches) | (ctrl, healthy, batch 2) | (ctrl, severe, batch 1) |
| Condition types 1&2 (w/ batch effect) | All | All | (stim, healthy, batch 2) | (ctrl, severe, batch 1) |

**Table S3.**
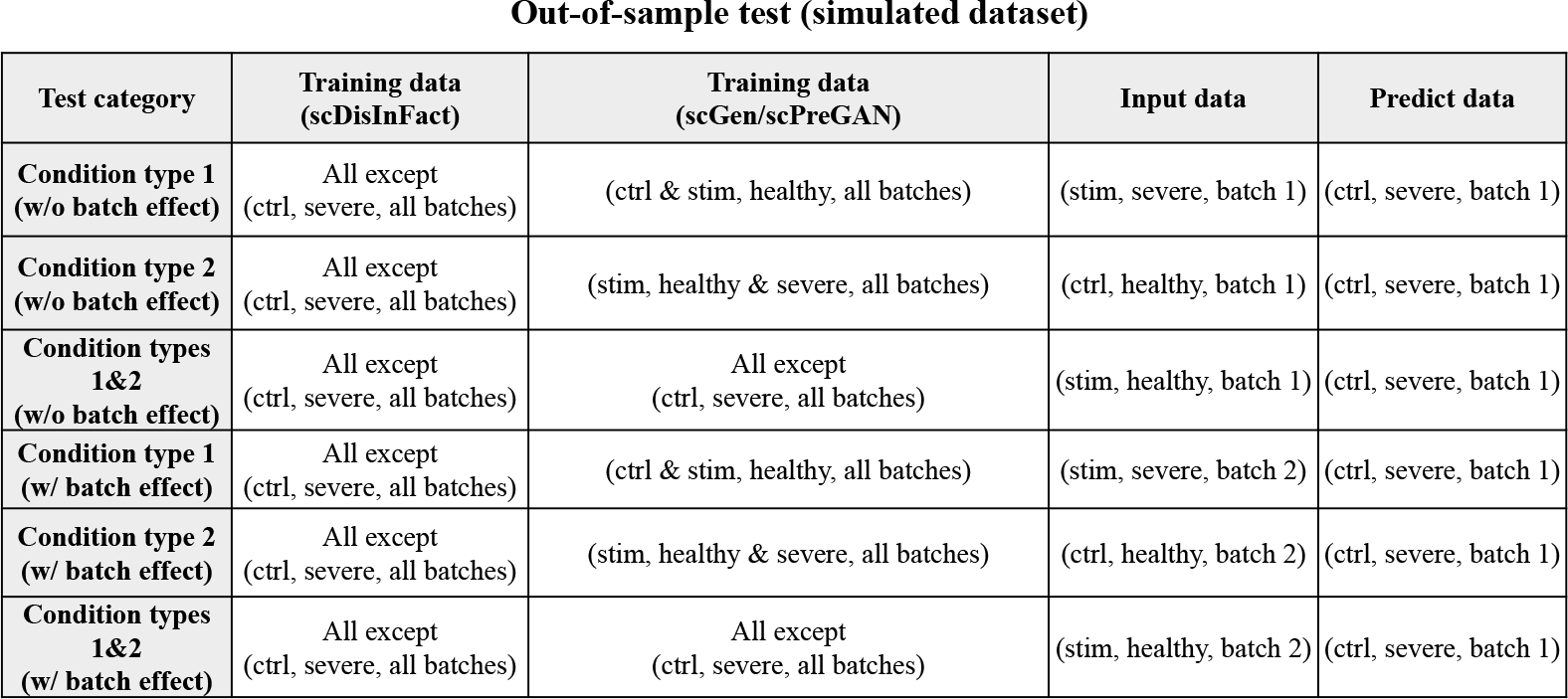
The setting of training data, input data, and predicted data of scDisInFact and baseline methods in *out-of-sample* prediction test on simulated datasets.

| Test category | Training data (scDisInFact) | Training data (scGen/scPreGAN) | Input data | Predict data |
| --- | --- | --- | --- | --- |
| Condition type 1 (w/o batch effect) | All except (ctrl, severe, all batches) | (ctrl & stim, healthy, all batches) | (stim, severe, batch 1) | (ctrl, severe, batch 1) |
| Condition type 2 (w/o batch effect) | All except (ctrl, severe, all batches) | (stim, healthy & severe, all batches) | (ctrl, healthy, batch 1) | (ctrl, severe, batch 1) |
| Condition types 1&2 (w/o batch effect) | All except (ctrl, severe, all batches) | All except (ctrl, severe, all batches) | (stim, healthy, batch 1) | (ctrl, severe, batch 1) |
| Condition type 1 (w/ batch effect) | All except (ctrl, severe, all batches) | (ctrl & stim, healthy, all batches) | (stim, severe, batch 2) | (ctrl, severe, batch 1) |
| Condition type 2 (w/ batch effect) | All except (ctrl, severe, all batches) | (stim, healthy & severe, all batches) | (ctrl, healthy, batch 2) | (ctrl, severe, batch 1) |
| Condition types 1&2 (w/ batch effect) | All except (ctrl, severe, all batches) | All except (ctrl, severe, all batches) | (stim, healthy, batch 2) | (ctrl, severe, batch 1) |

**Table S4.** The setting of prediction test when removing different numbers of count matrices in the training dataset.

| Test category | Training data |
| --- | --- |
| Hold out 1 matrix | All except<br>(ctrl, severe, batch 1) |
| Hold out 2 matrices | All except<br>(ctrl, severe, batch 1),<br>(stim, healthy, batch 2) |
| Hold out 3 matrices | All except<br>(ctrl, severe, batch 1),<br>(stim, healthy, batch 2),<br>(stim, severe, batch 1) |
| Hold out 4 matrices | All except<br>(ctrl, severe, batch 1),<br>(stim, healthy, batch 2),<br>(stim, severe, batch 1),<br>(ctrl, healthy, batch 2) |

**Table S5.** The patient ID, sample ID and condition labels of samples in GBM dataset

| Patient ID (Batch ID) | Sample ID | Condition |
| --- | --- | --- |
| PW029 | PW029-701 | Vehicle (DMSO) |
| PW030 | PW030-701 | Vehicle (DMSO) |
| PW030 | PW030-702 | Vehicle (DMSO) |
| PW030 | PW030-704 | 0.2 uM panobinostat |
| PW032 | PW032-702 | Vehicle (DMSO) |
| PW032 | PW032-705 | Vehicle (DMSO) |
| PW032 | PW032-709 | Vehicle (DMSO) |
| PW032 | PW032-711 | 0.2 uM panobinostat |
| PW034 | PW034-701 | Vehicle (DMSO) |
| PW034 | PW034-702 | Vehicle (DMSO) |
| PW034 | PW034-705 | 0.2 uM panobinostat |
| PW036 | PW036-701 | Vehicle (DMSO) |
| PW036 | PW036-702 | Vehicle (DMSO) |
| PW036 | PW036-705 | 0.2 uM panobinostat |
| PW040 | PW040-701 | Vehicle (DMSO) |
| PW040 | PW040-703 | Vehicle (DMSO) |
| PW040 | PW040-705 | Vehicle (DMSO) |
| PW040 | PW040-707 | Vehicle (DMSO) |
| PW040 | PW040-709 | Vehicle (DMSO) |
| PW040 | PW040-711 | Vehicle (DMSO) |
| PW040 | PW040-712 | 0.2 uM panobinostat |

**Table S6.** The detailed setting of training data, input data, and predicted data in perturbation prediction test on GBM dataset.

| Test category | Training data | Input data | Predict data |
| --- | --- | --- | --- |
| **Treatment**<br>**(w/o batch effect)** | All (except PW034-705) | (vehicle (DMSO), PW034) | (0.2 uM Panobinostat, PW034) |
| **Treatment**<br>**(w/ batch effect)** | All (except PW034-705) | (vehicle (DMSO), PW030) | (0.2 uM Panobinostat, PW034) |

**Table S7.** The detailed setting of training data, input data, and predicted data of scDisInFact and baseline methods in *in-sample* tests on the COVID-19 dataset.

| Test category | Training data<br>(scDisInFact) | Training data<br>(scGen/scPreGAN) | Input data | Predicted data |
| --- | --- | --- | --- | --- |
| **Age**<br>**(w/o batch effect)** | All | (moderate, 40-, all batches),<br>(moderate, 40-65, all batches) | (moderate, 40-, Lee_2020) | (moderate, 40-65, Lee_2020) |
| **Disease severity**<br>**(w/o batch effect)** | All | (healthy, 40-65, all batches),<br>(moderate, 40-65, all batches) | (healthy, 40-65, Lee_2020) | (moderate, 40-65, Lee_2020) |
| **Age & Disease severity**<br>**(w/o batch effect)** | All | (healthy, 40-65, all batches),<br>(healthy, 65+, all batches),<br>(moderate, 40-65, all batches) | (healthy, 65+, Lee_2020) | (moderate, 40-65, Lee_2020) |
| **Age**<br>**(w/ batch effect)** | All | (moderate, 40-, all batches),<br>(moderate, 40-65, all batches) | (moderate, 40-, Wilk_2020) | (moderate, 40-65, Lee_2020) |
| **Disease severity**<br>**(w/ batch effect)** | All | (healthy, 40-65, all batches),<br>(moderate, 40-65, all batches) | (healthy, 40-65, Wilk_2020) | (moderate, 40-65, Lee_2020) |
| **Age & Disease severity**<br>**(w/ batch effect)** | All | (healthy, 40-65, all batches),<br>(healthy, 65+, all batches),<br>(moderate, 40-65, all batches) | (healthy, 65+, Arunachalam_2020) | (moderate, 40-65, Lee_2020) |

**Table S8.**
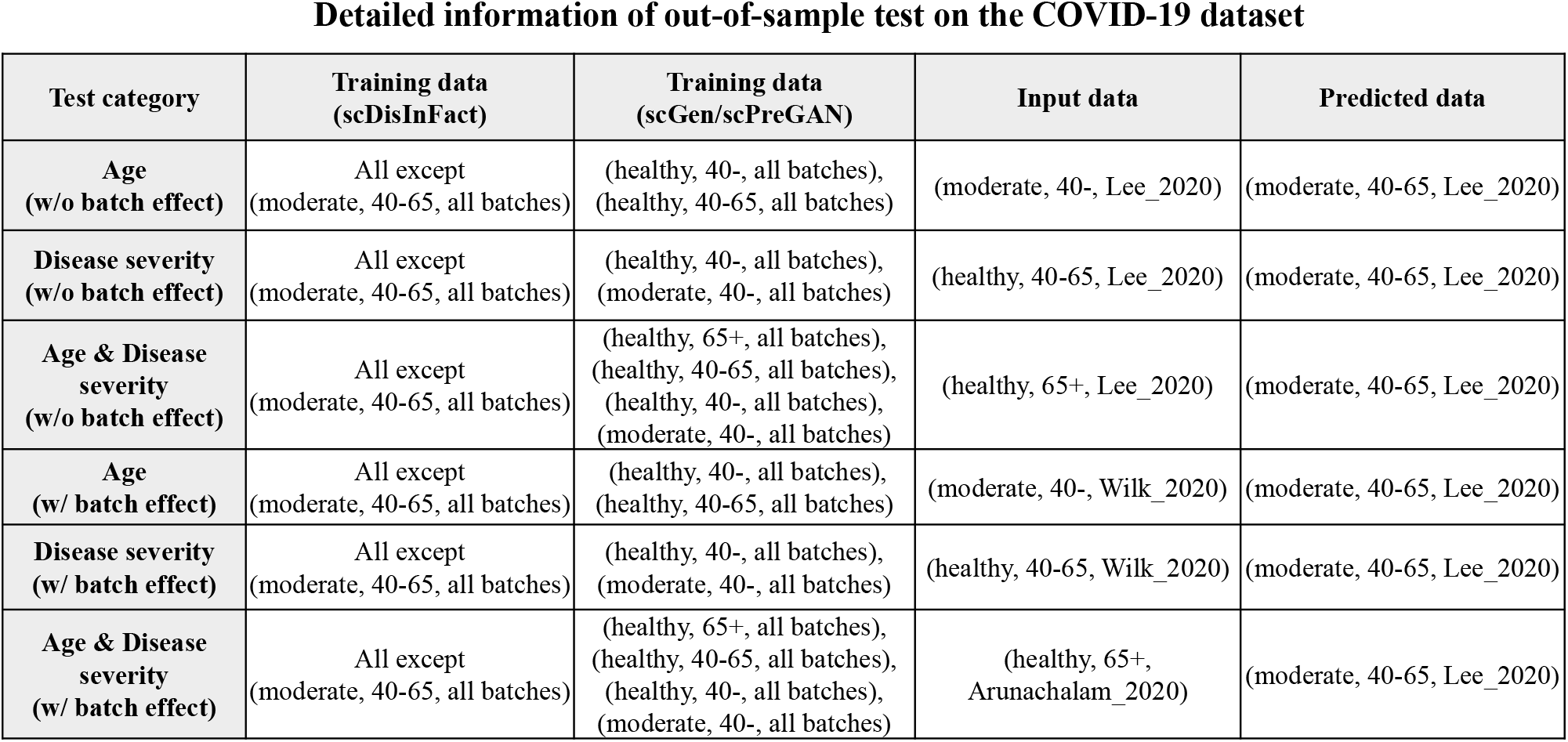
The detailed setting of training data, input data, and predict data of scDisInFact and baseline methods in *out-of-sample* test on the COVID-19 dataset.

| Test category | Training data<br>(scDisInFact) | Training data<br>(scGen/scPreGAN) | Input data | Predicted data |
| --- | --- | --- | --- | --- |
| **Age**<br>**(w/o batch effect)** | All except<br>(moderate, 40-65, all batches) | (healthy, 40-, all batches),<br>(healthy, 40-65, all batches) | (moderate, 40-, Lee_2020) | (moderate, 40-65, Lee_2020) |
| **Disease severity**<br>**(w/o batch effect)** | All except<br>(moderate, 40-65, all batches) | (healthy, 40-, all batches),<br>(moderate, 40-, all batches) | (healthy, 40-65, Lee_2020) | (moderate, 40-65, Lee_2020) |
| **Age & Disease**<br>**severity**<br>**(w/o batch effect)** | All except<br>(moderate, 40-65, all batches) | (healthy, 65+, all batches),<br>(healthy, 40-65, all batches),<br>(healthy, 40-, all batches),<br>(moderate, 40-, all batches) | (healthy, 65+, Lee_2020) | (moderate, 40-65, Lee_2020) |
| **Age**<br>**(w/ batch effect)** | All except<br>(moderate, 40-65, all batches) | (healthy, 40-, all batches),<br>(healthy, 40-65, all batches) | (moderate, 40-, Wilk_2020) | (moderate, 40-65, Lee_2020) |
| **Disease severity**<br>**(w/ batch effect)** | All except<br>(moderate, 40-65, all batches) | (healthy, 40-, all batches),<br>(moderate, 40-, all batches) | (healthy, 40-65, Wilk_2020) | (moderate, 40-65, Lee_2020) |
| **Age & Disease**<br>**severity**<br>**(w/ batch effect)** | All except<br>(moderate, 40-65, all batches) | (healthy, 65+, all batches),<br>(healthy, 40-65, all batches),<br>(healthy, 40-, all batches),<br>(moderate, 40-, all batches) | (healthy, 65+,<br>Arunachalam_2020) | (moderate, 40-65, Lee_2020) |

## Supplementary Note 1: Circle contrastive loss

Given the unshared-bio factors z*_u_* of a sample cell, we assume that there are *K* cells that have the same condition as the sample cell, and *L* cells that have different conditions. We then calculate the sample cell’s within-condition cosine similarity scores (denoted as 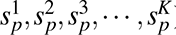), and cross-conditions cosine similarity scores (denoted as 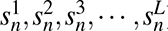). The circle contrastive loss on the cell maximizes its within-condition similarity while minimizing its cross-condition similarity:

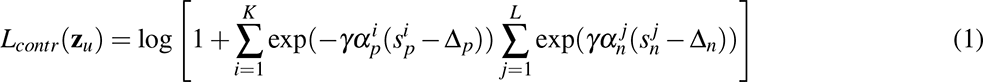

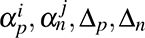 are decided by relaxation factor *m*^1^, which is set to be 0.25. *γ* is set to be 80.

## Supplementary Note 2: Maximum mean discrepancy loss

Given the shared-bio factors z_s_ from two different batches and conditions (denoted separately as 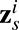 and 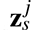), the MMD loss can be written as

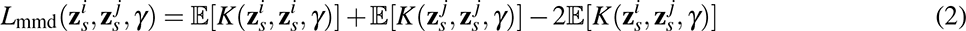

where *K*(*·,·, γ*) is a Gaussian kernel function of the form 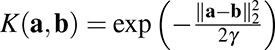, and *γ* is the hyper-parameter of the kernel function. Similar to^2^, we calculated MMD under different *γ*s to improve the robustness of the loss term

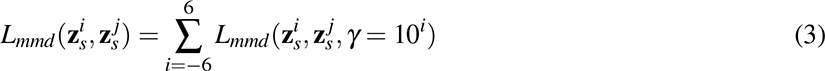

The same procedure also applies to unshared-bio factors z*_u_*.

